# Virome analyses by next-generation sequencing (NGS) in chilli (*Capsicum anuum* L.) presented with diverse symptoms phenotype revealed the association of seven plant viruses

**DOI:** 10.1101/2023.01.11.523546

**Authors:** Netla Vamsidhar Reddy, Shridhar Hiremath, Mantesh Muttappagol, H. D. Vinay Kumar, S. Koti Prasanna, T. L Mohan Kumar, C. R. Jahir Basha, K. S. Shankarappa, V. Venkataravanappa, C. N. Lakshminarayana Reddy

**Author notes:** Corresponding author’s mailing address: 1. Dr. C. N. Lakshminarayana Reddy, Department of Plant Pathology, College of Agriculture, University of Agricultural Sciences, Bengaluru, Karnataka, India.

## Abstract

Chilli is an important vegetable and spice crop, is known to be infected by several viruses. Techniques used in diagnosis of plant viral diseases before Next-generation sequencing (NGS) are having limitation of identifying the only known viruses. In the present study, virome analyses in infected chilli leaf samples was carried out using NGS to know the diversity of both known and unknown viruses associated with diseased symptoms. For virome profiling, samples from 19 fields were collected from chilli plants showing leaf curling, vein banding, mosaic, mottling, shoestring/rat tail/filiform/leathery and dull coloured leaves. Viral disease incidence in the surveyed fields varied from 26.66% to 47.50%. Total RNA was extracted from the 19 chilli leaf samples collected from fields and were pooled at equimolar concentration for virome profiling. From the rRNA-depleted pooled total RNA, mRNA and sRNA libraries were prepared and sequenced using Illumina NOVASEQ 6000 platform. Raw sequence data obtained was *de novo* assembled using three approaches; mRNAome with Trinity, sRNAome with Velvet and whole transcriptome (WT) with SPAdes assembly. Chilli virome, pairwise sequence identity and phylogenetic analyses revealed the presence of seven different viruses; chilli leaf curl virus (ChiLCV) along with its associated alpha and betasatellites, cucumber mosaic virus (CMV), groundnut bud necrosis orthotospovirus (GBNV), pepper cryptic virus-2 (PCV-2), pepper vein yellows virus (PeVYV), bell pepper alphaendornavirus (BPEV) and tobacco vein clearing virus (TVCV). From the virus associated contigs, complete/near-complete genomes for ChiLCV, CMV, PCV-2, PeVYV and BPEV and, partial genomes for GBNV and TVCV were reconstructed from the RNAome. Recombination breakpoint analyses revealed presence of recombination breakpoints in ChiLCV (coat protein and AC4 regions), CMV RNA2 (2a protein region) and P0, P3 and P5 protein regions of PeVYV. viruses identified in the current study are known to be originated from intra and interspecific recombination. Further, all the viruses detected in the pooled RNA sample were validated by PCR and loop mediated isothermal amplification (LAMP) using specific primers designed. Among the seven viruses identified in the study in chilli, PeVYV and BPEV are the first reports from India.

## INTRODUCTION

Chilli (*Capsicum annuum* L.) is a popular ethnopharmacological spice and vegetable crop grown throughout the world’s tropical and subtropical regions (Palukaitis et al., 1992). Due to its pungency, nutritional and economic importance, it is extensively cultivated throughout the year in countries like India (Khan et al., 2014). Chilli crop is vulnerable to a variety of DNA and RNA plant virus infections, and the number of viruses infecting chilli has increased dramatically over the last decade (Kenyon et al., 2014). Diverse viruses reported to infect chilli belongs to begomoviruses, potyviruses, cucumoviruses, tospoviruses, poleroviruses, criniviruses, alfamoviruses, tobamoviruses, tymoviruses, ilarviruses, tombusviruses, cryptoviruses, nepoviruses and endornaviruses. These viruses are mainly transmitted by whitefly, aphids, thrips and nematodes in nature. In addition to this, some them are also known to have mechanical and seed transmission (Kenyon et al., 2014). Viruses infecting chilli produce varied symptoms such as, curling, mosaic, stunting and mottling, and these can vary significantly with cultivar, age of the plant, strain of the virus and environmental conditions (Waweru et al., 2019). In the recent decade, there is an increase in outbreaks of virus species infecting chilli, which has become a major constraint to its production. Hence, early detection and diagnosis of viruses infecting chilli and its carrier vectors, play a crucial role in reducing disease spread and developing effective management strategies (Hema and Konakalla, 2021).

However, many viruses have gone unnoticed or are yet to be identified in chilli. Under field conditions, mixed infections will be encountered, which results in diverse symptoms manifestations. Through the detection methods like enzyme-linked immunosorbent assay (ELISA), polymerase chain reaction (PCR), and their variants, only expected or known viruses can be detected, leaving the status of all other unknown viruses in the infected samples (Czotter et al., 2018). The drawbacks of these methodologies can be overcome by studying the virome profile (total viral population) through next-generation sequencing (NGS) technologies (Sidharthan et al., 2020).

NGS enables the direct detection, identification and discovery of viruses in an unbiased manner without requiring any antibodies or prior knowledge of the pathogen macromolecular sequence (Barzon et al., 2011). NGS, combined with informatics for *de novo* discovery and assembly of plant virus or viroid genome has been used since 2009, for discovering novel DNA and RNA viruses (Adams et al., 2009; Kreuze et al., 2009). Based on the NGS technology, the novel viruses infecting many crops (peaches, lilies, barley, sweet pepper, grapevine, vanilla, and citrus) have been detected in different parts of the world (Pecman et al., 2017; Jo et al., 2018a; Jo et al., 2018b; Wright et al., 2020).

Among the various NGS based approaches for virome analyses, mRNA and sRNA sequencing emerged as the most reliable approaches for detection of plant viruses in recent times (Pirovano et al., 2015; Maliogka et al., 2018; Pooggin, 2018; Massart et al., 2019; Sidharthan et al., 2020). From mRNA and sRNA libraries, single-stranded (ss) as well as double-stranded (ds) RNA viruses and DNA viruses can be effectively captured (Seguin et al., 2014; Roossinck et al., 2015; Jo et al., 2017). Virome analyses by NGS identify viruses in a broad spectrum and accurately, allowing for multiplex detection, quantification, and the discovery of new viruses. Apart from this, the technique is likely to become the future standard for diagnostics as its cost decreases and becomes more affordable (Rubio et al., 2020).

Considering the importance of plant viruses in chilli crop production in India, an attempt was made to virome profile through NGS in chilli samples collected from chilli growing areas of Karnataka, India. In the current manuscript, results of known and novel viruses detected in chilli samples, their genome construction and development of diagnostics are discussed.

## MATERIAL AND METHODS

### Survey for virus disease incidence and collection of virus infected samples for virome profiling

Survey was conducted for the incidence of viral diseases in chilli in major chilli growing areas of Bengaluru (Mavallipura and Muthagadahalli) and Kolar (Rajakallahalli, Madderi and Mallandahalli) districts of Karnataka, India. Percent disease incidence of viral diseases was calculated (Senanayake et al., 2012) and symptoms on the infected plants were recorded (Figure 1). Nineteen chilli leaf samples from infected plants exhibiting different symptoms were collected for virome profiling and detailed information regarding the collected samples has been given in the supplementary table 1. Leaf samples were snap-freezed with liquid nitrogen in the field and preserved with proper labelling and stored in a deep freezer (−80°C) until further processing. Leaf samples were ground into a fine powder using liquid nitrogen and used for RNA and DNA extraction.

**Figure 1.**
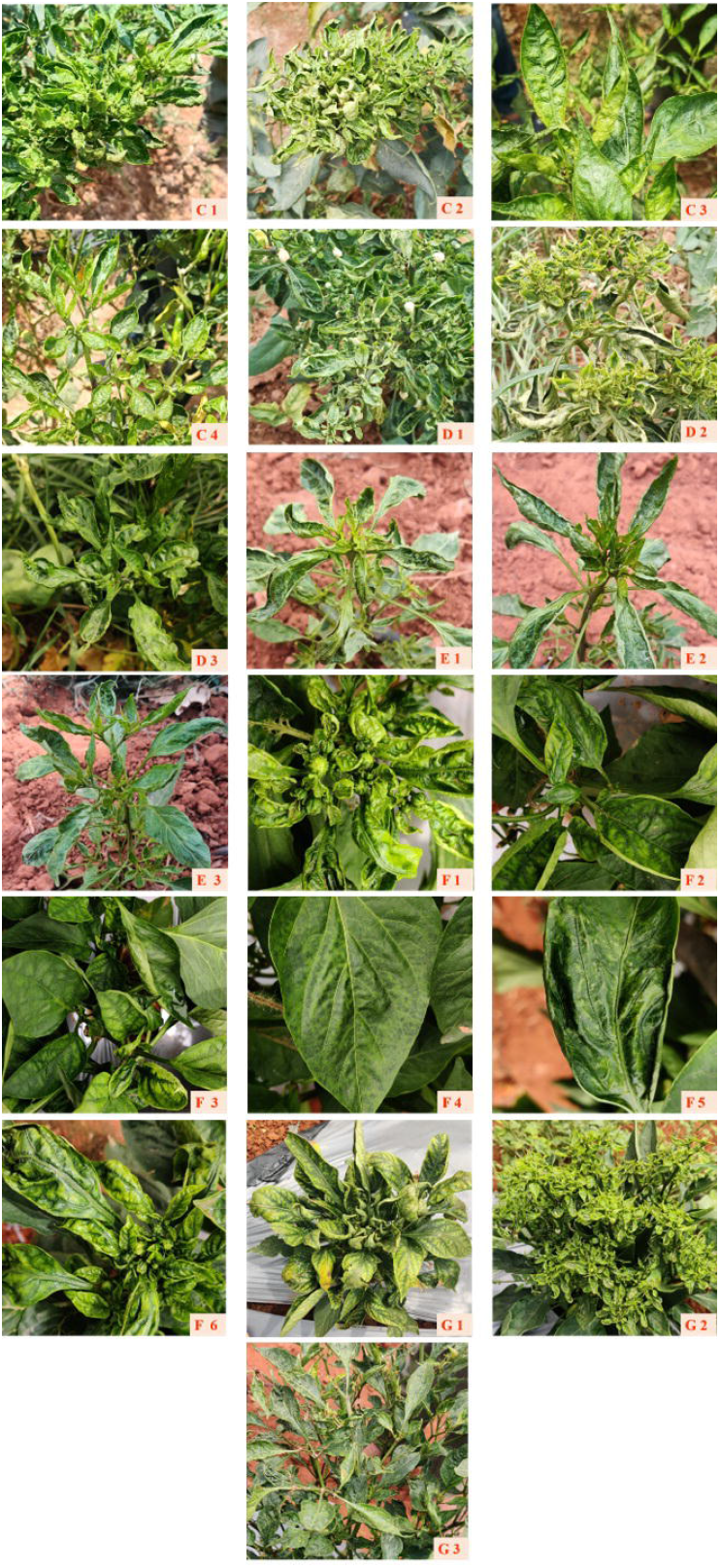
Chilli plants showing different viral disease symptoms under natural conditions, from which samples were collected for virome profiling. Symptoms observed in each sample are as follows. C1) leaf curling and mosaic, C2) leaf curling, mosaic and bunchy appearance of leaves, C3) mottling and dark green vein banding, C4) leathery leaves and vein banding, D1) leaf curling, D2) leaf curling and mosaic, D3) mottling and mosaic of leaves, E1) leaf curling and mosaic, E2) mosaic and rat tailed leaves, E3) mosaic, F1) leaf curling, mottling and vein banding, F2) mottling and vein banding, F3) vein banding, F4) mosaic, F5) mottling and mosaic, F6) mottling and vein banding, G1) leaf curling and vein banding, G2) leaf curling and small leaves and G3) mottling, mosaic and dull coloured leaves.

### RNA extraction, library construction and next-generation sequencing

Total RNA was extracted from nineteen leaf samples by phenol-chloroform and lithium-chloride (LiCl) method (Sajeevan et al., 2014; Khairul-Anuar et al., 2019) and then quantified using Nanodrop (Thermo Fisher Scientific, USA) and Agilent 4150 Tapestation system for purity and integrity, respectively. Total RNA purity and integrity analysis was done by bleach agarose gel electrophoresis according to procedure outlined by Aranda et al. (2012). Good quality total RNA extracted from nineteen leaf samples was mixed at equimolar concentrations and created a pool of total RNA samples. From the pooled total RNA sample, ribosomal RNA was removed by using the Ribo-Zero rRNA Removal Kit (Illumina, San Diego, CA, USA). From the rRNA depleted total RNA, mRNA and sRNA libraries were constructed by Illumina trueseq stranded mRNA library preparation kit (Cat. No. 20020594) and TruSeq small RNA sample preparation guide (Illumina, San Diego, CA, United States) as per the manufacturer’s protocol, respectively. The libraries were quantified using Qubit 4.0 fluorometer (Thermofisher #Q33238) using a DNA HS assay kit (Thermofisher #Q32851) following manufacturer’s protocol. To identify the insert size of the library, we queried it on Tapestation 4150 (Agilent) utilizing highly sensitive D1000 screen tapes (Agilent # 5067-5582) following the manufacturer’s protocol. Libraries were sequenced on Illumina NOVASEQ 6000 platform facility available at Neuberg Center for Genomic Medicine, Neuberg Supertech reference laboratories, Ahmedabad, to generate 150-bp paired-end reads.

### Raw data pre-processing

Raw mRNA and sRNA NGS data quality were checked by FastQC Version 0.11.9 (FASTQ Quality Check) tool (http://www.Bioinformatics.babraham.ac.uk/projects/fastqc) (Andrews, 2010). Sequence adapters and low-quality reads were trimmed by Trim Galore version 0.6.5 with a quality Phred score of q=20 (https://www.bioinformatics.babraham.ac.uk/projects/trim_galore) (Krueger, 2015). After trimming, the quality of the reads was once again checked with FastQC.

### *De-novo* assembly

Good quality mRNA and sRNA cleaned reads were used for *de novo* assembly. Three approaches were followed for *de novo* assembly to better unravel the chilli virome. The mRNA data was assembled using Trinity version 2.13.2 (Grabherr et al., 2011) (https://github.Com/trinityrnaseq/releases/tag/Trinity-v2.13.2) with default parameters, sRNA data was assembled using Velvet version 1.2.10 (Zerbino and Birney, 2008) and whole transcriptome (WT) (combined mRNA and sRNA data) data was *de novo* assembled using SPAdes version 3.13.1 (Bushmanova et al., 2019)

### Identification of viruses

Assembled contigs generated through *de novo* assembly from three approaches were subjected to standalone MEGABLAST analysis (e-value cut off: Ie^-5^; query coverage: ≥90%) against the complete reference sequences of viruses available (https://www.ncbi.nlm.nih.gov/genome/viruses/) at NCBI GenBank (National centre for biotechnology information) blast + version 2.12.0 (Camacho et al., 2009) (https://ftp.ncbi.nlm.nih.Gov/blast/executables/blast+/LATEST/). Contigs with length >200 (for mRNA and WT) and >50 (for sRNA) nucleotides (nt) were considered for analyses. All the identified viruses were categorized based on the ICTV guidelines (International committee on taxonomy of viruses) and threshold levels of nt identity given for viral species and strains demarcations. The ICTV threshold levels for viral species and strain demarcation are listed in supplementary table 2.

### Reconstruction of genome of identified viruses

Virus associated contigs were filtered from total contigs using Sequence alignment/map (SAM) tools version 1.9 (Li et al., 2009). Identified viral contigs were aligned against reference virus isolates retrieved from the GenBank to construct complete/ near-complete viral genome using the Clustal X version 2.0 (Larkin et al., 2007) (https://clustalx.software.informer.com/2.1/) and BioEdit version 7.2 (Hall et al., 2011) (https://bioedit.software.informer.com/). From the reconstructed complete/near complete viral genomes, all of the anticipated open reading frames (ORFs) were recovered for identified viruses using NCBI ORFfinder (https://www.ncbi.nlm.nih.gov/orffinder/).

### Sequence comparison

Complete genomes retrieved from NCBI GenBank along with the viral genomes reconstructed in this study were aligned using Clustal W, which is integrated in Sequence Demarcation Tool (SDT) version 1.2 (Muhire et al., 2014) to calculate the pairwise sequence identity scores for comparison of reconstructed genomes of identified viruses with their respective genomes retrieved from NCBI.

### Phylogenetic analysis

Complete genomes retrieved from NCBI GenBank along with the viral genomes reconstructed in this study were aligned using Clustal W tool, which is integrated in MEGA 11 programme (Tamura et al., 2021). Aligned sequences were subjected to phylogenetic tree construction using Neighbourhood Joining (NJ) method with the selection of model Kimura 2-parameter (K2P) model with 1000 bootstrap replicates in MEGA 11 programme.

### Recombination analysis

To know the possible recombination events in identified viruses from the study, by using Clustal W aligned MEGA file as input, recombination analysis was performed using RDP5 (Martin et al., 2021) package integrated with six different algorithms (Recombination Detection Program (RDP), GENECONV, Max Chi, Chimaera, Si Scan and 3Seq). Recombination events detected by at least three algorithms in the reconstructed viral genomes were considered significant. A Set of virus sequences used for phylogenetic analysis were utilized for recombination analysis.

Overall work flow of virome analyses in chilli is represented in figure 2 as flow chart.

**Figure 2.**
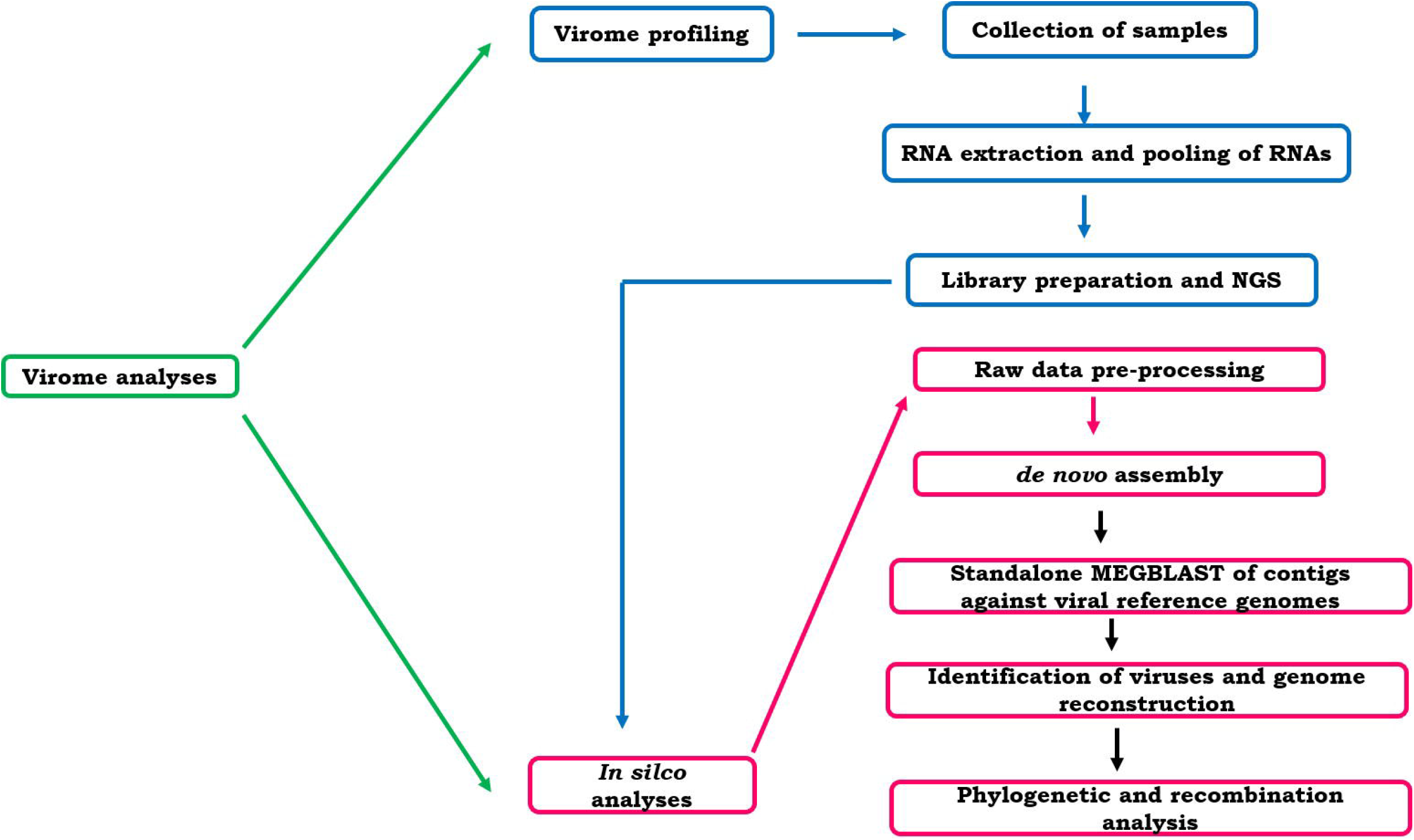
Virome analyses work flow in chilli. Chilli leaf samples showing varied disease symptoms were collected for virome profiling. Total RNA extracted from each sample was pooled and used for mRNA and sRNA libraries preparation. Libraries of mRNA and sRNA were sequenced through next-generation sequencing (NGS) platforms and obtained raw data was pre-processed. Good quality mRNA and sRNA data was *de novo* assembled and obtained assembled contigs were used for identification and genome reconstruction of the viruses. Reconstructed complete/near-complete viral genomes were used for phylogenetic and recombination analyses.

### Validation of identified viruses

Chilli virome analyses identified both DNA and RNA viruses. Presence of DNA virus was validated by polymerase chain reaction (PCR) and loop-mediated isothermal amplification (LAMP). Similarly, presence of RNA viruses was validated by reverse-transcription PCR (RT-PCR) and reverse-transcription LAMP (RT-LAMP). Specific primers were designed targeting the coat protein (ChiLCV, CMV, PCV-2 and PeVYV), movement protein (GBNV) and poly protein (BPEV) genes. Details of the expected amplicons length for the primers designed for each virus are listed in table 1A and 1B. PCR and LAMP were performed using pooled total DNA sample prepared from pooling of DNA extracted from 19 chilli leaf samples. RT-PCR and RT-LAMP assays were performed with cDNA (complementary DNA) prepared from pooled total RNA. Synthesis of c DNA from 1000 ng of pooled total RNA was carried out by using 10 μM of reverse primer from each reconstructed RNA virus and RevertAid Reverse Transcriptase (Thermo Fisher scientific, USA) (Cat. No. EP0441) according to the manufacturer’s protocol.

**Table 1A:**
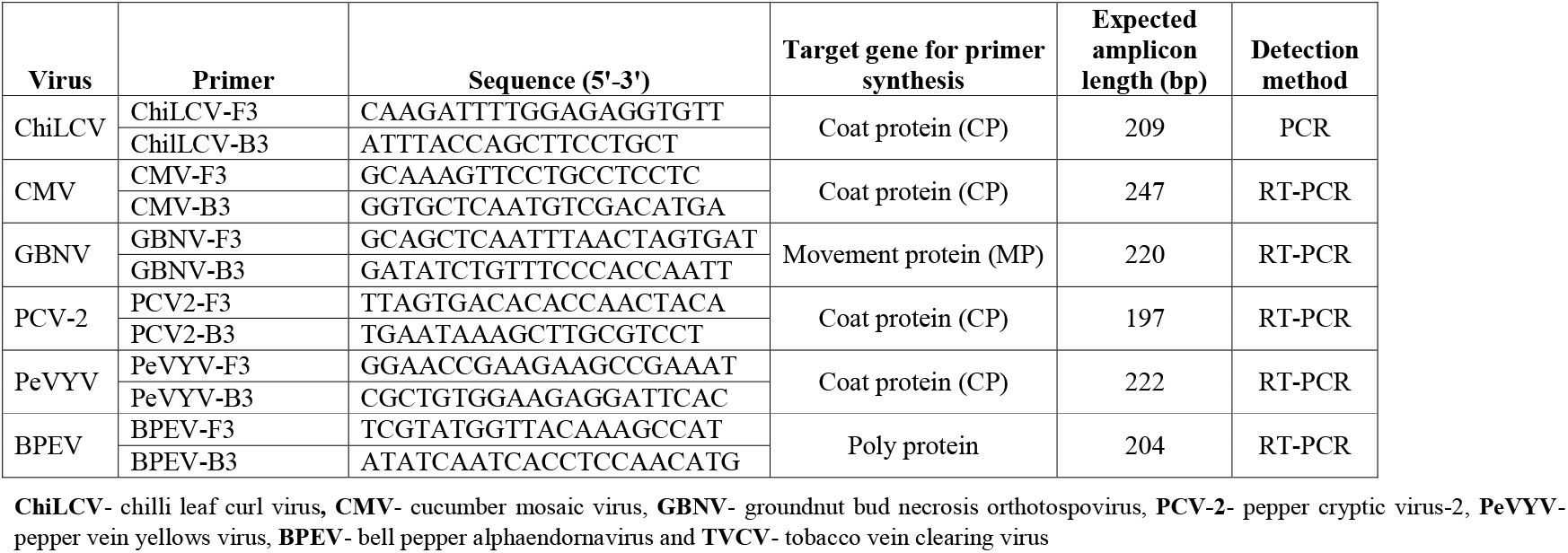
PCR and RT-PCR primers designed for identified viruses from the virome analyses for detection and validation

**Table 1B:**
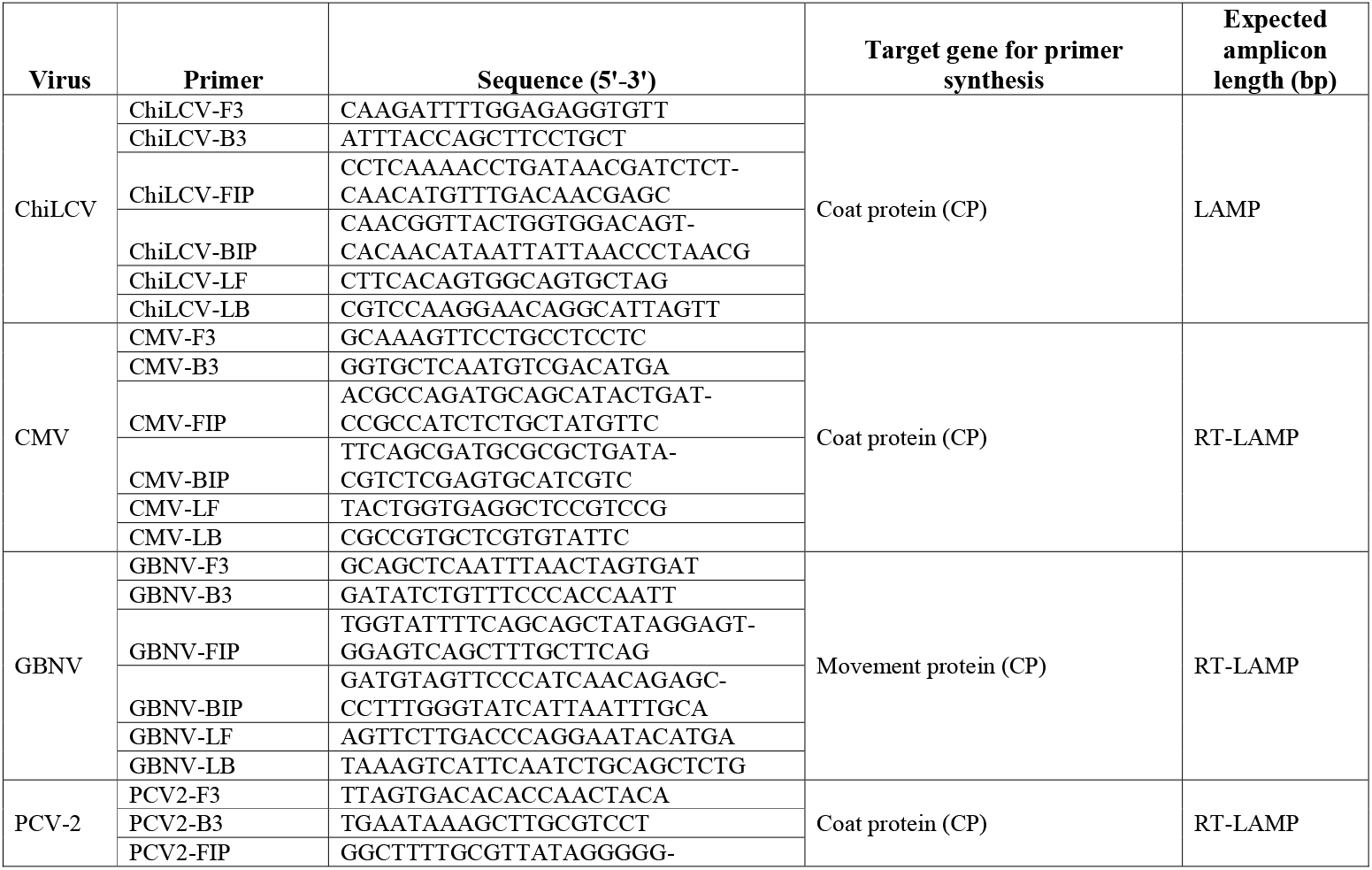
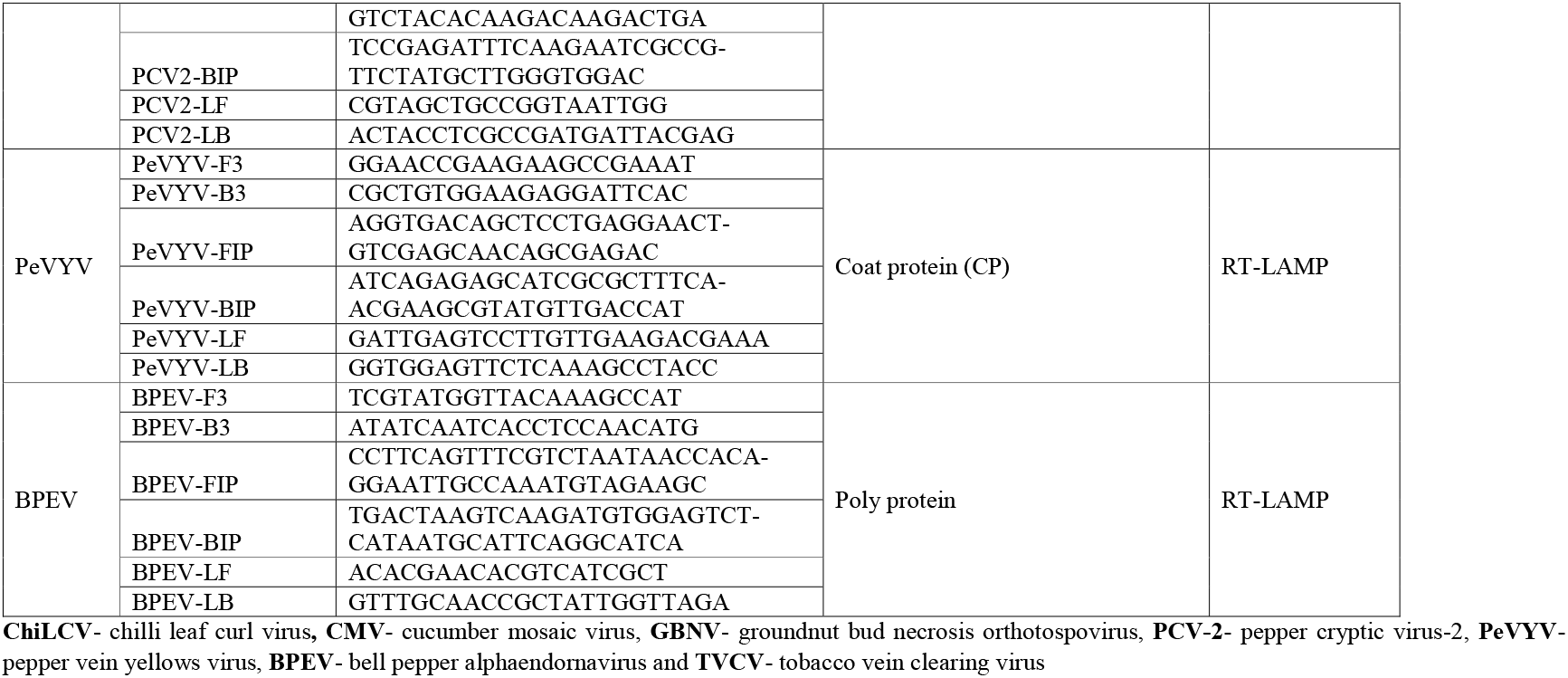
LAMP and RT-LAMP primers designed for identified viruses from the virome analyses for detection and validation

### PCR and LAMP assays

PCR assay was performed with initial denaturation of 94° C for 5 minutes, 35 cycles of denaturation for 30 seconds at 94° C, annealing for 30 seconds at 50° C, extension for 45 sec at 72° C and final extension of 72° C for 10 minutes for all the identified viruses. PCR reactions were carried out in ProFlex PCR system (Applied biosystems, Thermo Fisher Scientific, USA). All the PCR amplifications were performed in a reaction volume of 20 μL PCR mix containing 2 μL DNA (50 ng/μL)/ 5 μL cDNA (20 ng/μL) template, 10 μL of EmeraldAmp^®^ GT PCR Master Mix (Takara Bio, USA), 10 μM of each forward and reverse primers. Amplification products were analysed on 1% agarose gel (80V for 1 hr) stained with ethidium bromide (10 mg/mL) and viewed on Gel Doc XR Imaging System (BioRad, USA).

LAMP assays were carried out with 25 μL LAMP mixture containing 2 μL of DNA/cDNA (50 ng/μL) template, 1 μL (10 μM) of F3, B3, 2 μL (10 μM) of FIP, BIP, LF and LB primers, 1.5 μL of 10 mM dNTPs, 1.0 μL of 5 M betaine, 6.5 μL of sterile double-distilled water, 2.5 μL of 1X ThermoPol Reaction buffer (20 mM Tris–HCl, 10 mM (NH_4_)SO_4_, 10 mM KCl, 2 mM MgSO_4_, 144 0.1% Triton X-100, pH 8.8 @ 25 °C), 0.5 μL of 100 mM MgSO4, and 1.0 μL of 8U *Bst* DNA Polymerase (New England Biologicals, USA). Reaction mixture was incubated in a hot water bath at 60°C for 1 hr followed by 80 °C for 10 min to terminate the reaction. LAMP reaction products were analysed on 2% agarose gels (65V for 3 hr) stained with ethidium bromide (10 mg/mL) and viewed on Gel Doc XR Imaging System (BioRad, USA).

For the visual detection of LAMP reaction products, nucleic acid staining dyes VeriPCR dye (Chromus Biotech PVT Ltd, Bengaluru, India) and hydroxy naphthol blue (HNB) (Sigma Aldrich, USA) were used. Five μL of 1X VeriPCR dye was added after amplification, whereas, 0.2 μL of 20 mM HNB was added to the LAMP master mix prior to amplification for visual inspection of the reaction products. Positive result with VeriPCR dye shows green fluorescent and HNB shows a colour change from violet to sky blue.

## RESULTS

### Viral disease incidence, symptomatology and samples preparation for virome profiling, NGS data and *de novo* assembly

Viral diseases were prevalent in all surveyed areas (Kolar and Bengaluru Districts) of Karnataka State, India with incidence ranging from 26.66% to 47.50%. Infected plants exhibited different symptoms such as leaf curling, vein banding, mosaic, mottling, shoestring/rat tail/filiform/leathery and dull coloured leaves (Figure 1). Total RNA from nineteen leaf samples collected from the infected chilli plants showing different symptoms was extracted and pooled in equimolar concentration to get a common RNA pool for virome profiling.

The mRNA and sRNA libraries prepared from the rRNA depleted pooled total RNA was used for NGS. The Illumina sequencing generated raw reads of 15.8 (5.70 GB) and 31.6 (4.70 GB) million sequences for mRNA and sRNA libraries, respectively. Raw sequencing of mRNA and sRNA data obtained were deposited in the SRA database with Bio project ID: PRJNA914908 accession numbers SRR22863580 and SRR22876797, respectively. The mRNA and sRNA reads (without adapter sequences, poor quality reads and Phred score> 20) were proceeded directly for *de novo* assembly. The *de novo* assembly of mRNA with Trinity, sRNA with Velvet and whole transcriptome (WT) with SPAdes software generated contigs of 38262, 8773 and 41383, respectively.

### Identification of viruses

Assembled contigs were subjected to MEGABLAST analysis against the complete reference sequences of viruses retrieved from NCBI GenBank revealed the presence of both DNA virus along with their associated satellites and RNA viruses. BLASTn search results of mRNA, sRNA and WT libraries showing the contigs associated with viruses are listed in supplementary tables 3, 4 and 5, respectively. From mRNA, sRNA and WT libraries, seven viruses were identified such as chilli leaf curl virus (ChiLCV), cucumber mosaic virus (CMV), groundnut bud necrosis orthotospovirus (GBNV), pepper cryptic virus-2 (PCV-2), pepper vein yellows virus (PeVYV), bell pepper alphaendornavirus (BPEV) and tobacco vein clearing virus (TVCV) (Supplementary table 6A). Identified viruses belonged to six different families such as *Geminiviridae* (ChiLCV), *Bromoviridae* (CMV), *Tospoviridae* (GBNV), *Partitiviridae* (PCV-2), *Solemoviridae* (PeVYV) and *Caulimoviridae* (TVCV) (Figure 3A). Identified ChiLCV from chilli virome was associated with tomato leaf curl Bangladesh betasatellite (ToLCBDB), tomato leaf curl Anand alphasatellite (ToLCAA), croton yellow vein mosaic alphasatellite (CYVMA), ageratum yellow vein Singapore alphasatellite (AYVSA) and tomato leaf curl Virudhunagar alphasatellite strain severe (ToLCVirA). Both mRNA and WT libraries revealed presence of seven viruses such as ChiLCV, CMV, PCV-2, PeVYV, BPEV and TVCV in the infected chilli samples. From sRNA library, five viruses (ChiLCV, CMV, PCV-2, PeVYV and BPEV) were identified. Among the DNA satellites of begomoviruses, ToLCBDB was identified in all three datasets, ToLCAA was identified in sRNA and WT library, CYVMA was identified in mRNA and sRNA library, AYVSA and ToLCVirA were identified only in sRNA library (Supplementary table 6A and 6B and figure 3B). According to our knowledge, PCV-2 is the first report in southern part of India, PeVYV and BPEV are the first reports from India on chilli.

**Figure 3.**
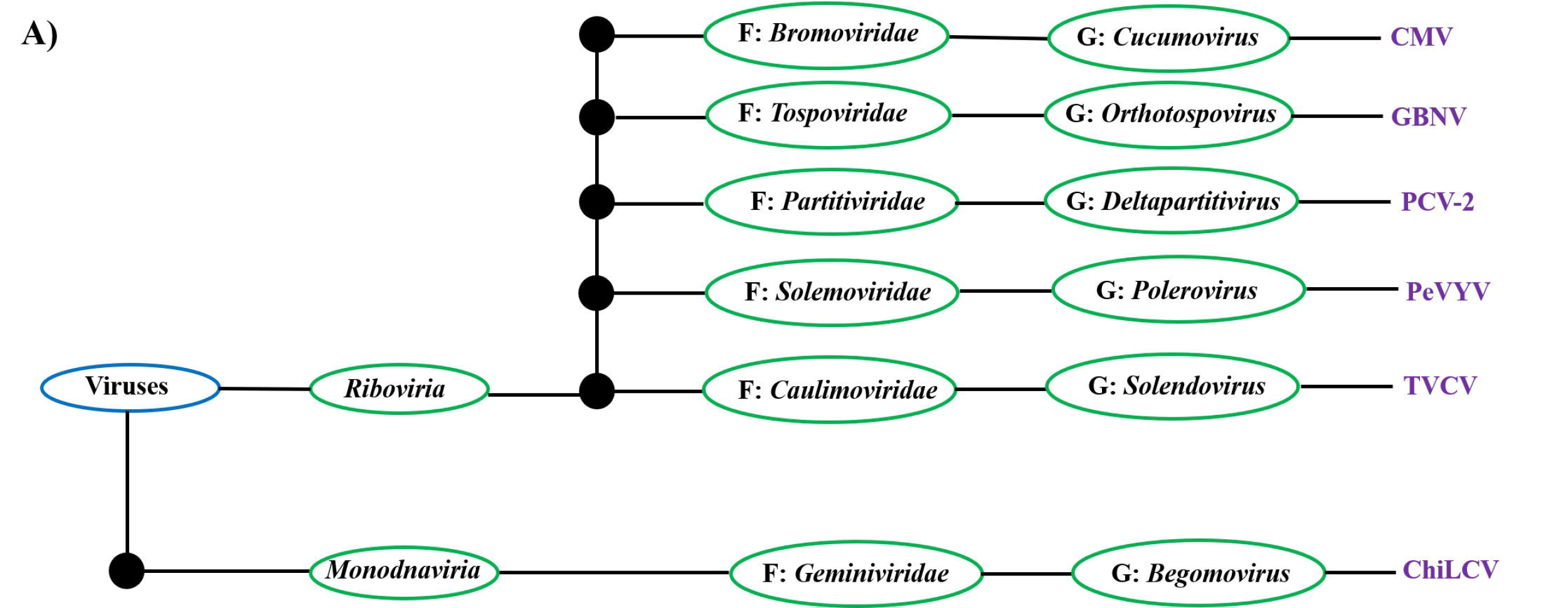
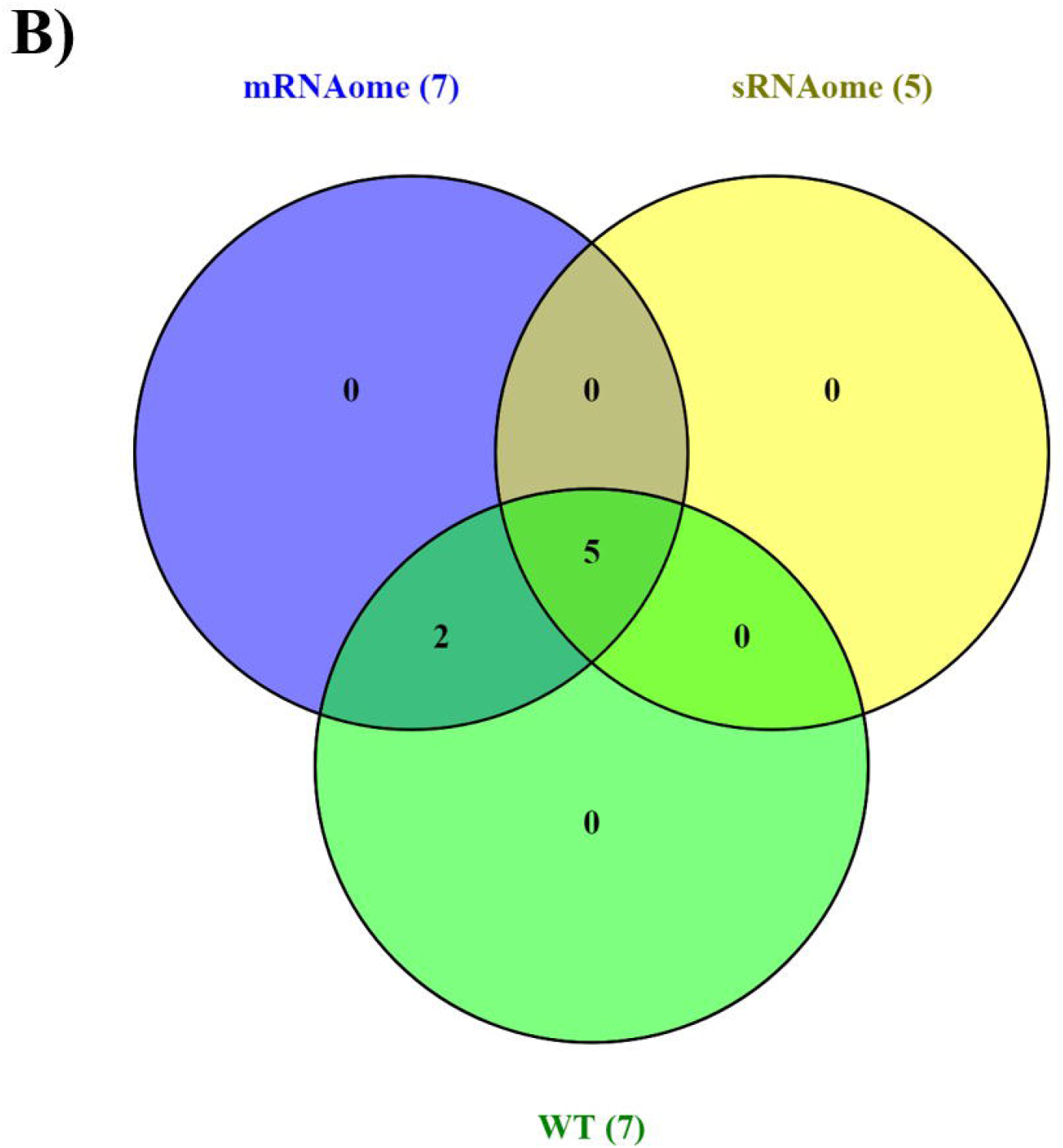
Identified viruses from chilli mRNA, sRNA and whole transcriptome (WT) libraries. A) Classification of identified viruses based on the ICTV taxonomy. B) Venn diagram showing the identified viruses in mRNAome (ChiLCV, CMV, GBNV, PCV-2, PeVYV, BPEV and TVCV), sRNAome (ChiLCV, CMV, PCV-2, PeVYV and BPEV and WT (ChiLCV, CMV, GBNV, PCV-2, PeVYV, BPEV and TVCV). Three different coloured circles represent the library name and number of identified viruses from the library.

### Reconstruction of genomes of identified viruses

Virus-associated contigs from the total contigs of mRNA, sRNA and WT libraries were separated and reconstructed into complete or near-complete genomes. Number of contigs associated with identified viruses from mRNA, sRNA and WT libraries have been listed in table 2 and shown in figure 4A. From the total contigs, by mapping virus-associated contigs to the NCBI designated reference genome of identified viruses we obtained complete or near-complete genomes (>90%) for five viruses (ChiLCV, CMV, PCV-2, PeVYV and BPEV). For two identified viruses (GBNV and TVCV), the viral contigs could not be assembled into complete genomes. Trinity assembly with mRNA and SPAdes assembly with WT libraries yielded longer contigs compared to Velvet assembly with sRNA library. For reconstruction of ChiLCV, CMV (RNA1, RNA2 and RNA3), PCV-2 (RNA1), GBNV (L and M segments), BPEV and PeVYV genomes, SPAdes assembled virus-associated contigs were used which resulted in reconstruction of genome with coverage of >90% complete genome except for GBNV L and M segments, which have genome coverage of 9.13 and 20.25 %, respectively (Table 2 and figure 4B). PeVYV and BPEV genomes were reconstructed with maximum coverage of 99.50 and 99.92 %, respectively. PCV-2 RNA2, GBNV S segment and TVCV genome were reconstructed using Trinity assembled with genome coverage of 97.70, 21.82 and 17.46 %, respectively (Table 2 and Figure 4B). All of the anticipated ORFs were identified from the complete or near complete reconstructed viruses with the aid of NCBI ORFfinder (Supplementary table 7). Genome organization of the complete reconstructed viruses are depicted in figure 4C-G. Full length intact ORFs identification was not done for the partially reconstructed genomes. Reconstructed viral genomes are deposited in the NCBI GenBank with the accession numbers listed in supplementary table 8. Complete/near-complete genomes of ChiLCV, CMV, PCV-2, PeVYV and BPEV were considered for further analyses.

**Figure 4.**
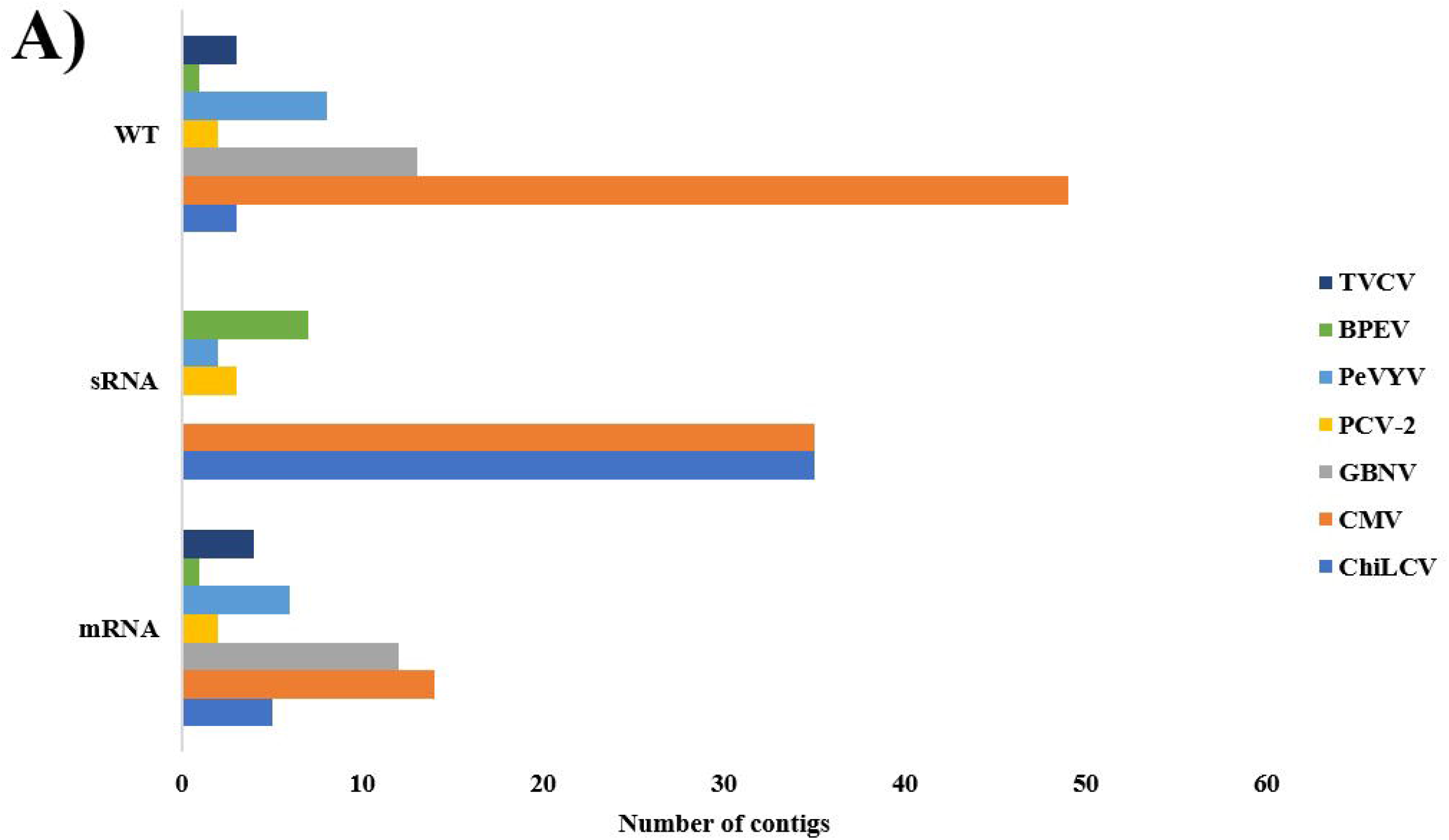
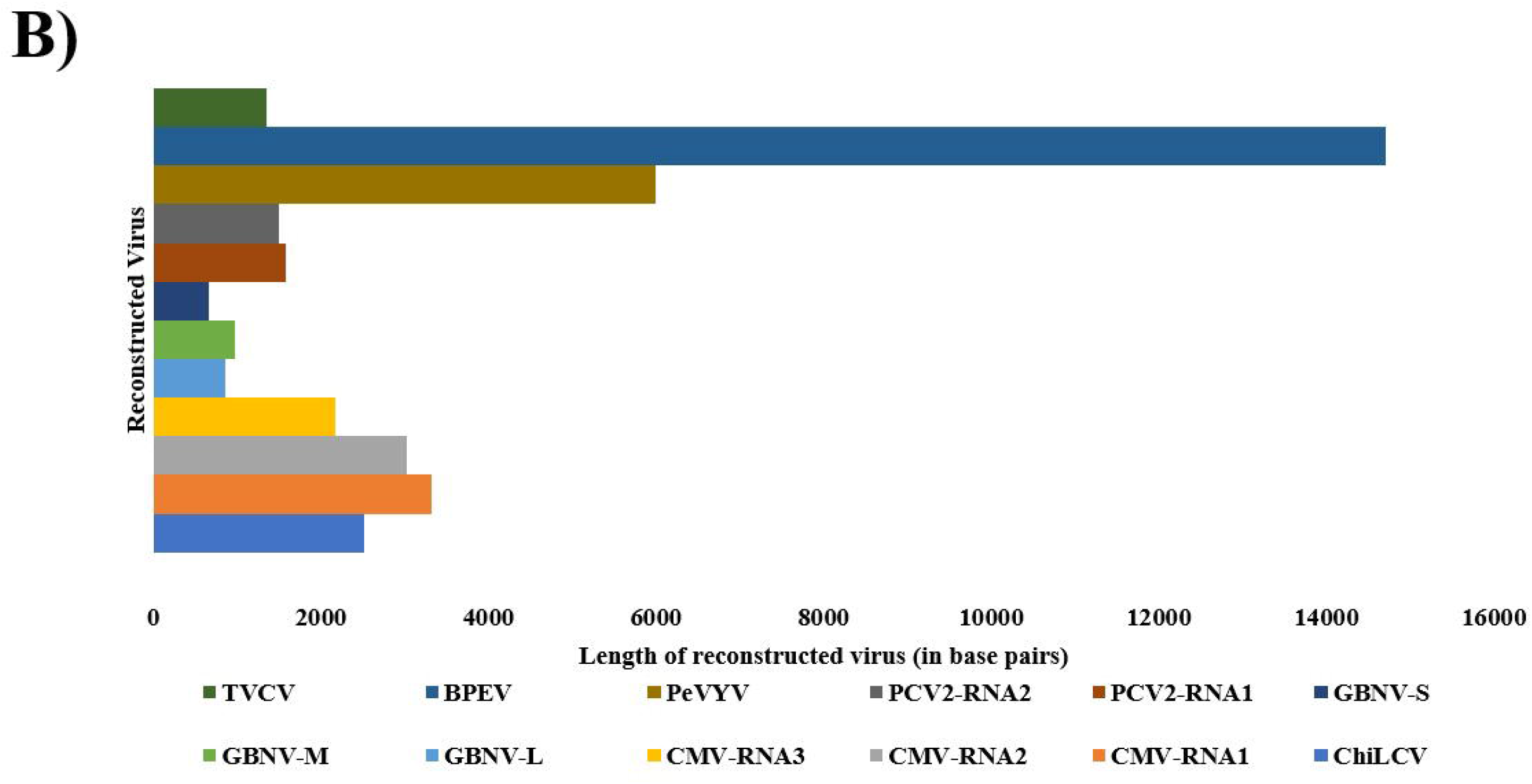
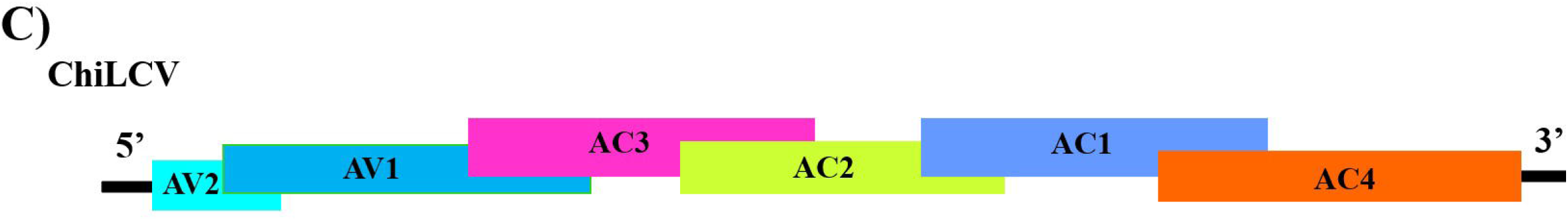
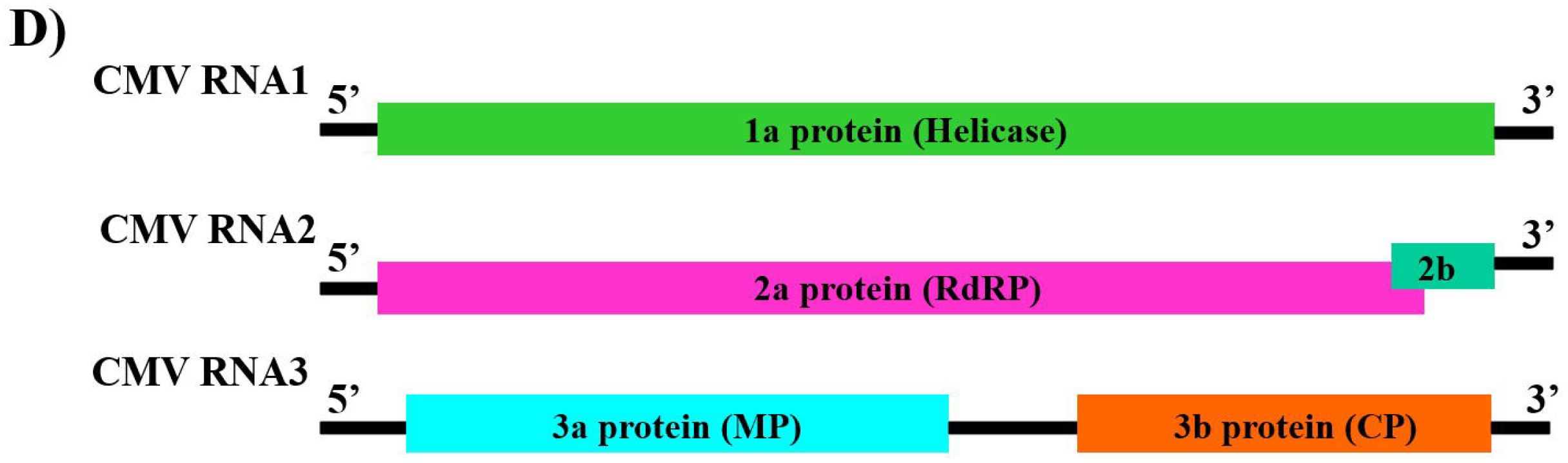
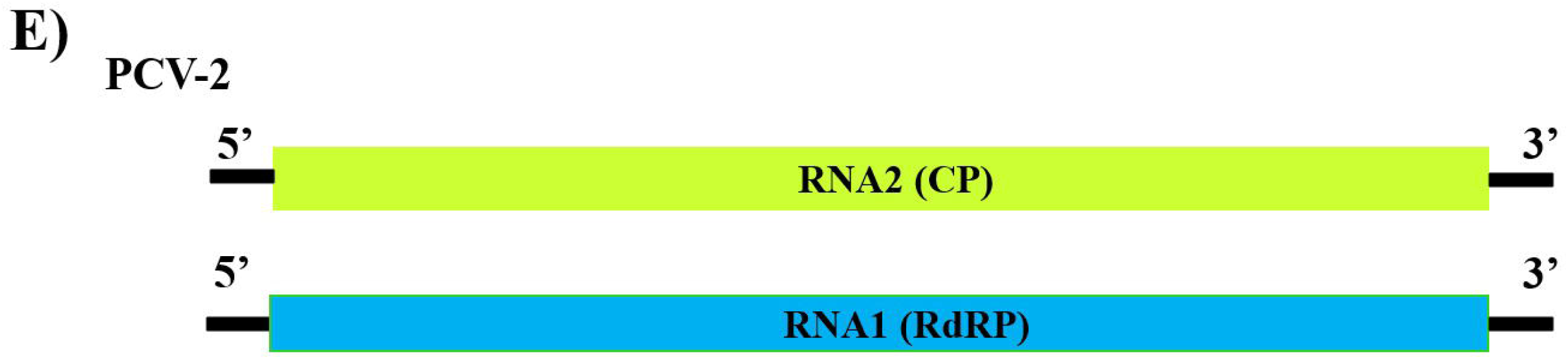
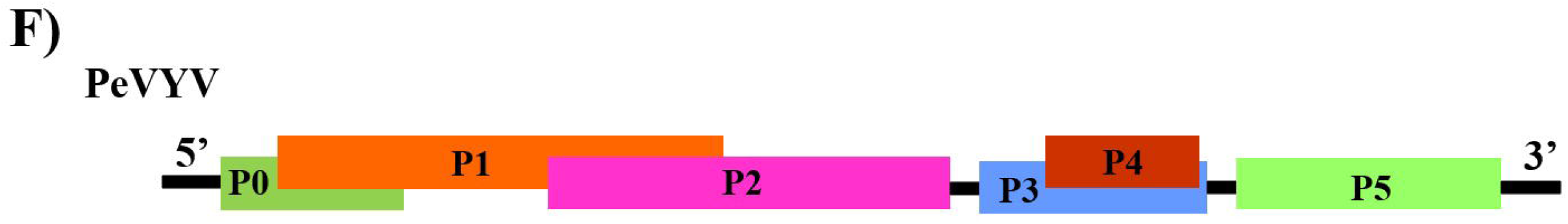
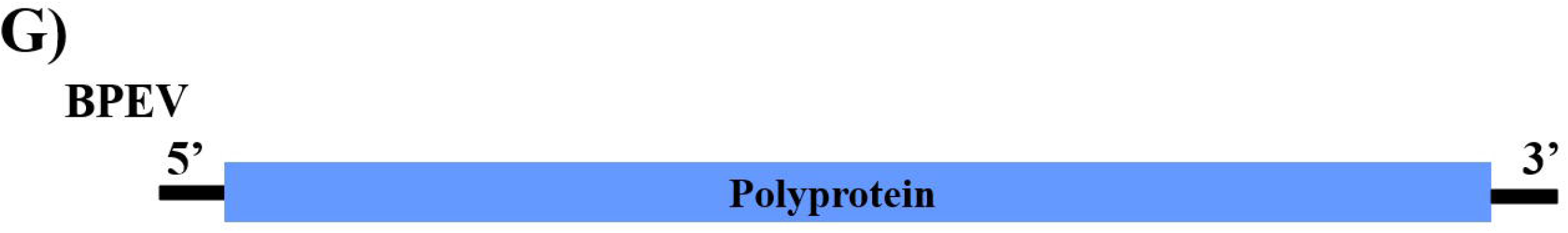
A) Graphical representation of number of contigs associated with each identified virus in mRNA, sRNA and WT library B) Graph showing the length of reconstructed viral genomes. Genome organization of reconstructed complete/near complete genomes C) ChiLCV, D) CMV, E) PCV-2, F) PeVYV and G) BPEV.

**Table 2:** Reconstructed viral genome sequences with their lineage and summary of virus-associated contigs in mRNA, sRNA and whole transcriptome (WT) library

| Genome sequence of the virus reconstructed | Segment | Genome | No. of contigs associated with virus |  |  | Reference genome | Size (nt) | Genome recovered (nt) | Percentage of genome recovery |
| --- | --- | --- | --- | --- | --- | --- | --- | --- | --- |
|  |  |  | mRNA | sRNA | WT |  |  |  |  |
| ChiLCV | DNA-A | Circular, ssDNA | 5 | 35 | 3 | NC_004628.1 | 2754 | 2510 | 91.14 |
| CMV | RNA1 | Linear, (+) ssRNA | 6 | 11 | 18 | NC_002034.1 | 3357 | 3313 | 98.69 |
|  | RNA2 |  | 3 | 13 | 7 | NC_002035.1 | 3050 | 3016 | 98.89 |
|  | RNA3 |  | 5 | 11 | 24 | NC_001440.1 | 2216 | 2175 | 98.15 |
| GBNV | L segment | Linear, (-) ssRNA | 9 | 0 | 6 | NC_003614.1 | 8911 | 859 | 9.64 |
|  | M segment |  | 3 | 0 | 5 | NC_003620.1 | 4801 | 972 | 20.25 |
|  | S segment |  | 0 | 0 | 2 | NC_003619.1 | 3057 | 667 | 21.82 |
| PCV-2 | RNA1 | Linear, dsRNA | 1 | 3 | 1 | NC_034159.1 | 1609 | 1585 | 98.51 |
|  | RNA2 |  | 1 | 0 | 1 | NC_034167.1 | 1525 | 1490 | 97.70 |
| PeVYV |  | Linear, (+) ssRNA | 6 | 2 | 8 | NC_055129.1 | 6028 | 5998 | 99.50 |
| BPEV |  | Linear, dsRNA | 1 | 7 | 1 | NC_039216.1 | 14714 | 14702 | 99.92 |
| TVCV |  | Circular, dsDNA | 4 | 0 | 3 | NC_003378.1 | 7767 | 1356 | 17.46 |

**ChiLCV**- chilli leaf curl virus, **CMV**- cucumber mosaic virus, **GBNV**- groundnut bud necrosis orthospovirus, **PCV-2**- pepper cryptic virus-2, **PeVYV**- pepper vein yellows virus, **BPEV**- bell pepper alphaendornavirus and **TVCV**- tobacco vein clearing virus

### Sequence comparison and phylogenetic analysis

Complete/near complete genomes (ChiLCV, CMV, PCV-2, PeVYV and BPEV) identified from chilli virome were subjected to sequence comparison and phylogenetic analyses (Figure 5A-H) along with the complete genomes of other viral sequences retrieved from the NCBI GenBank. Pairwise identity scores of identified viruses with NCBI retrieved genomes are listed in supplementary table 9-17).

**Figure 5.**
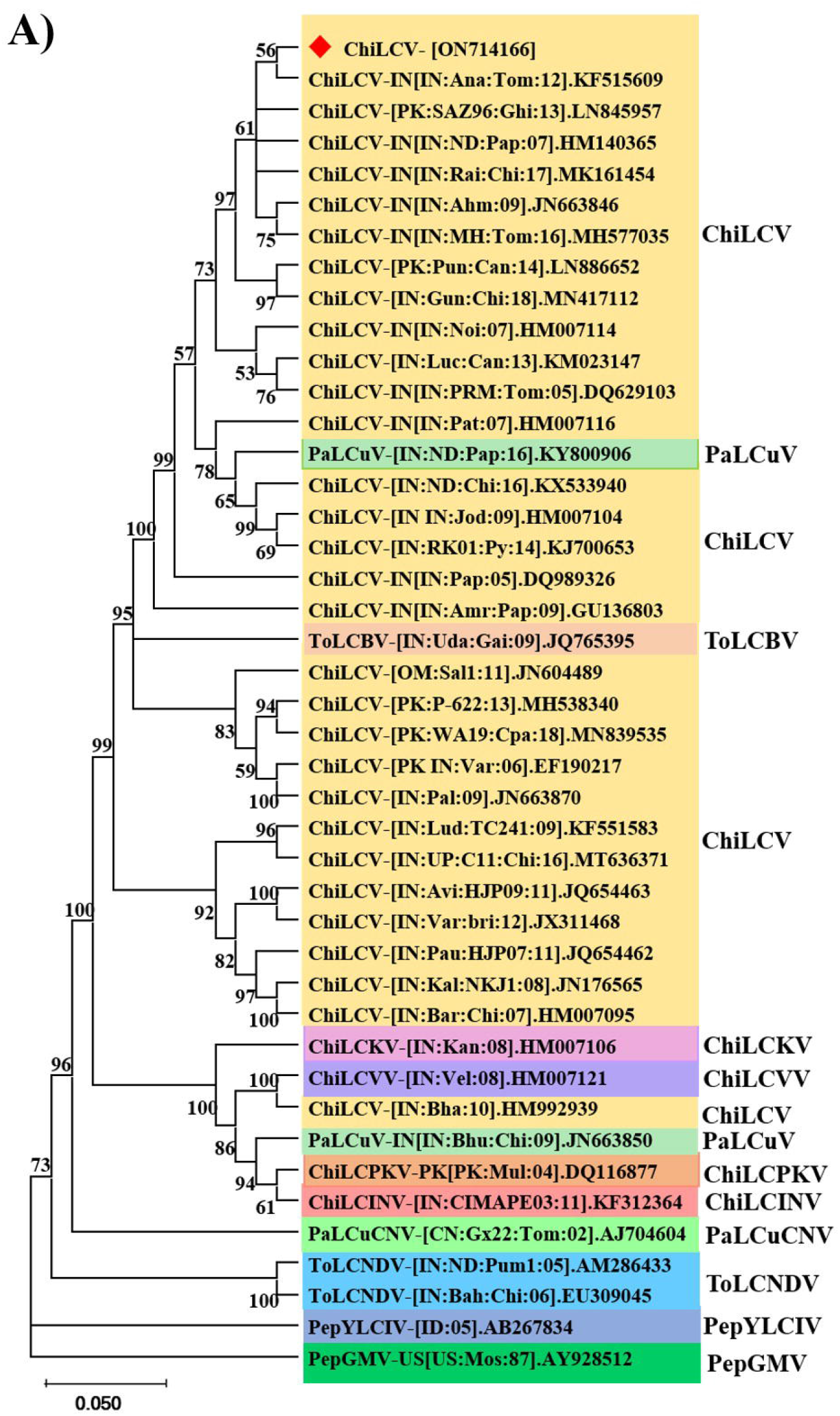
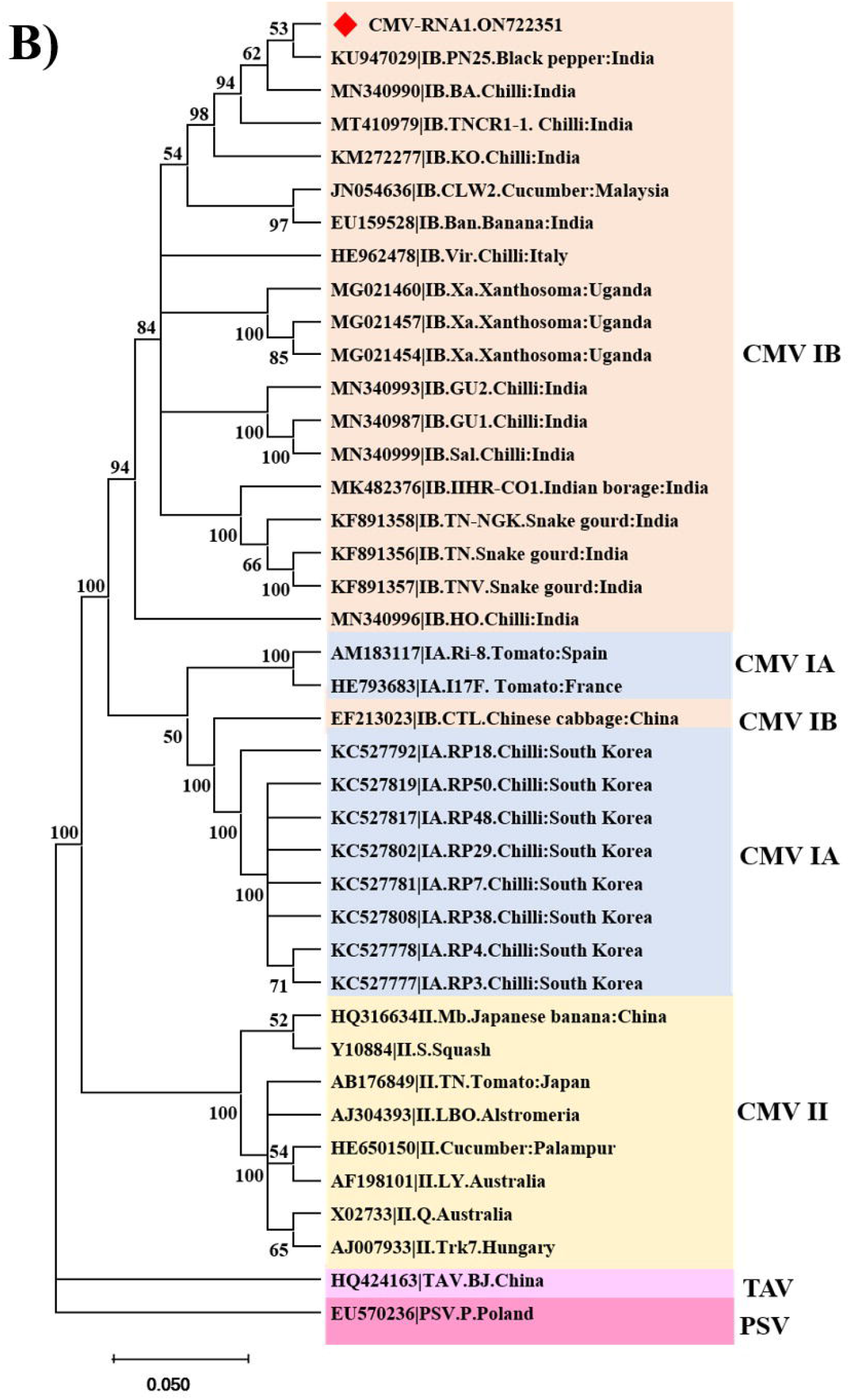
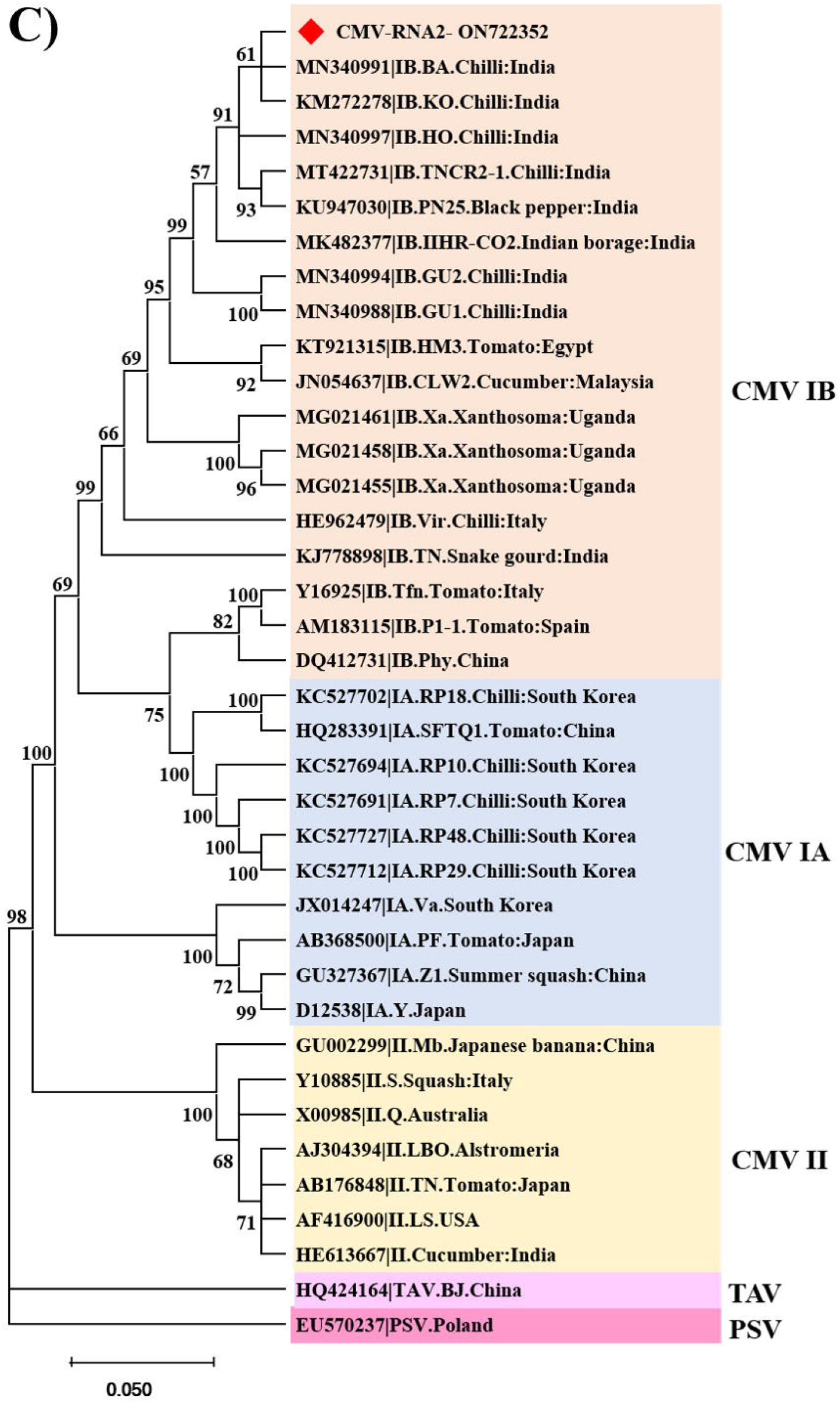
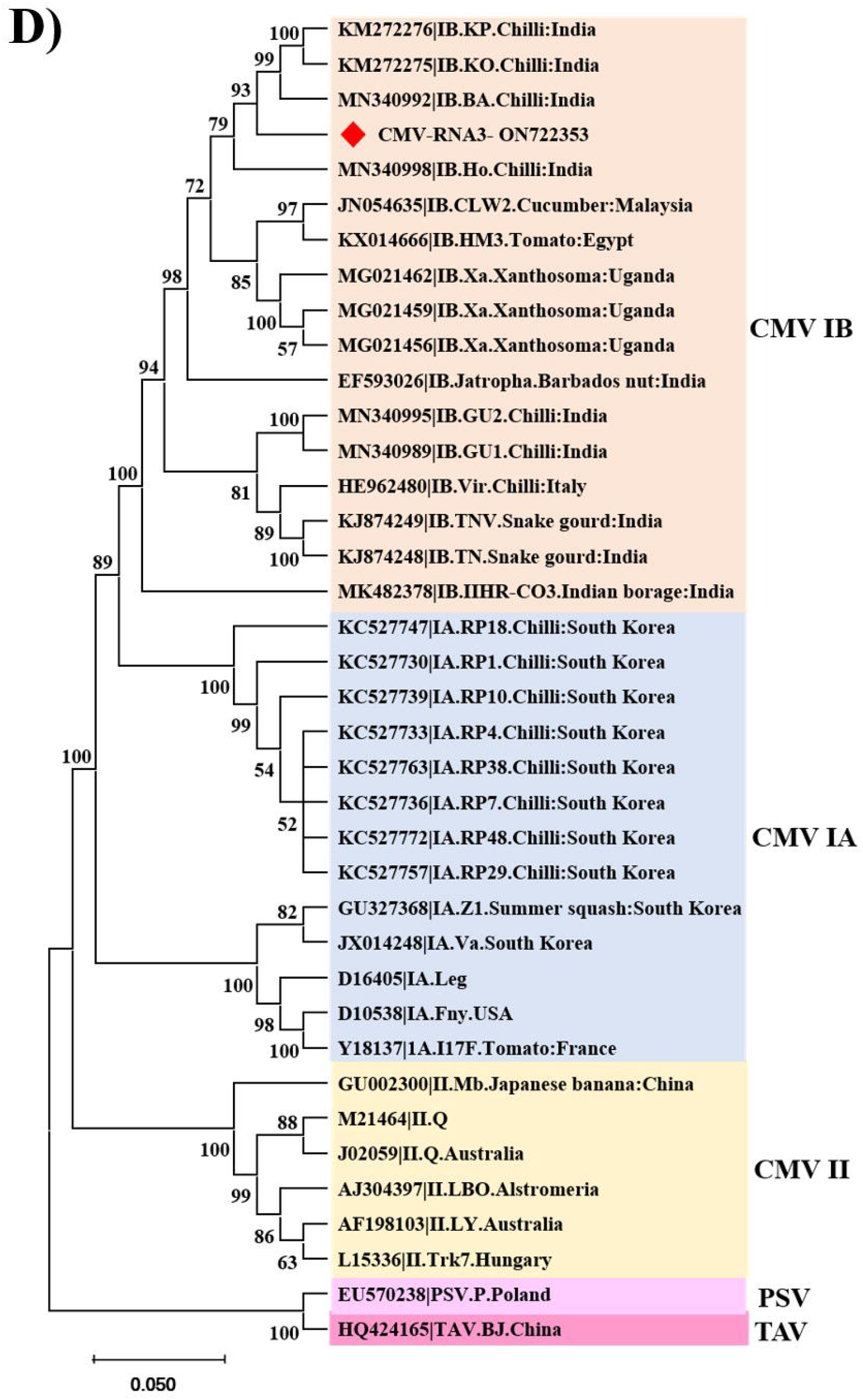
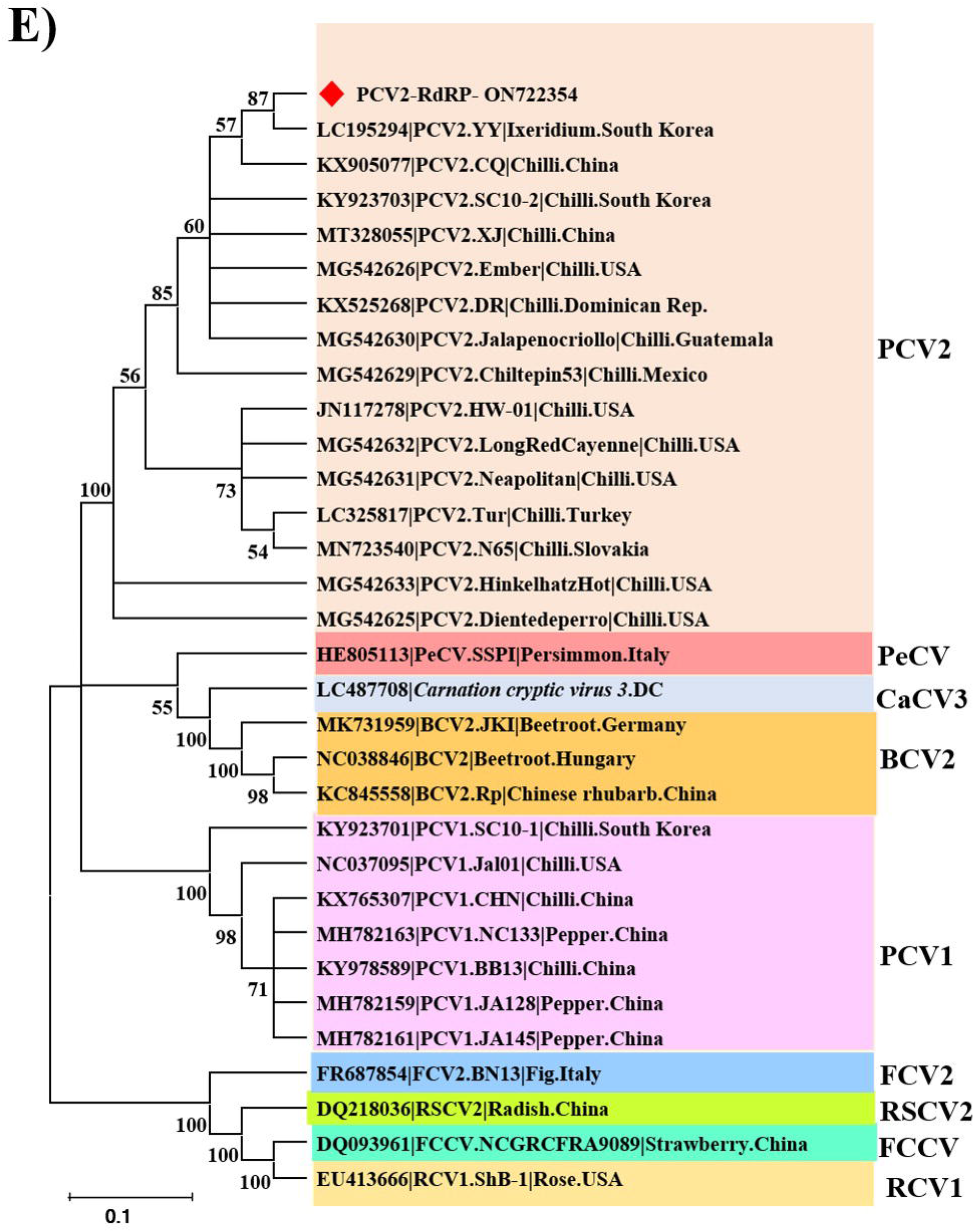
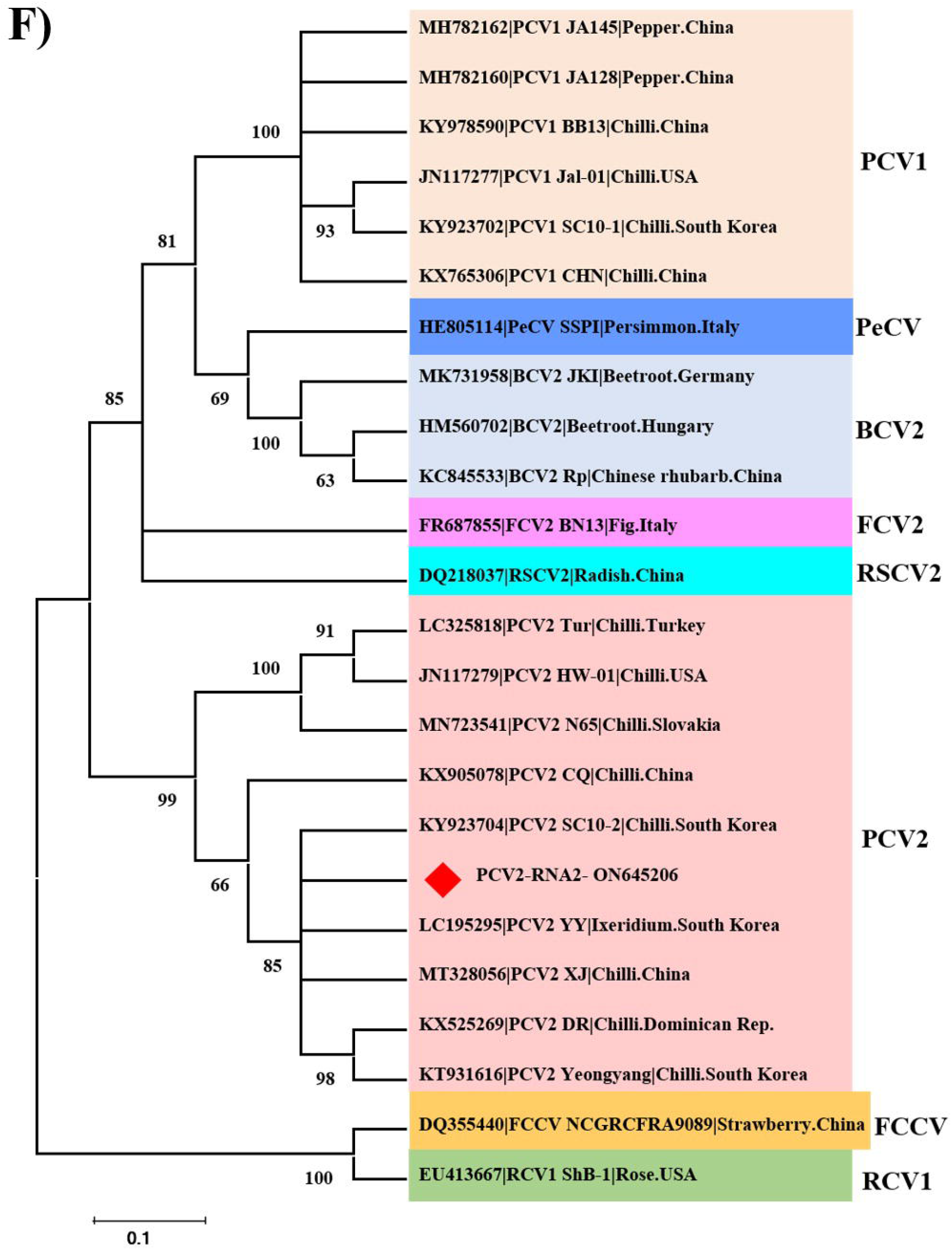
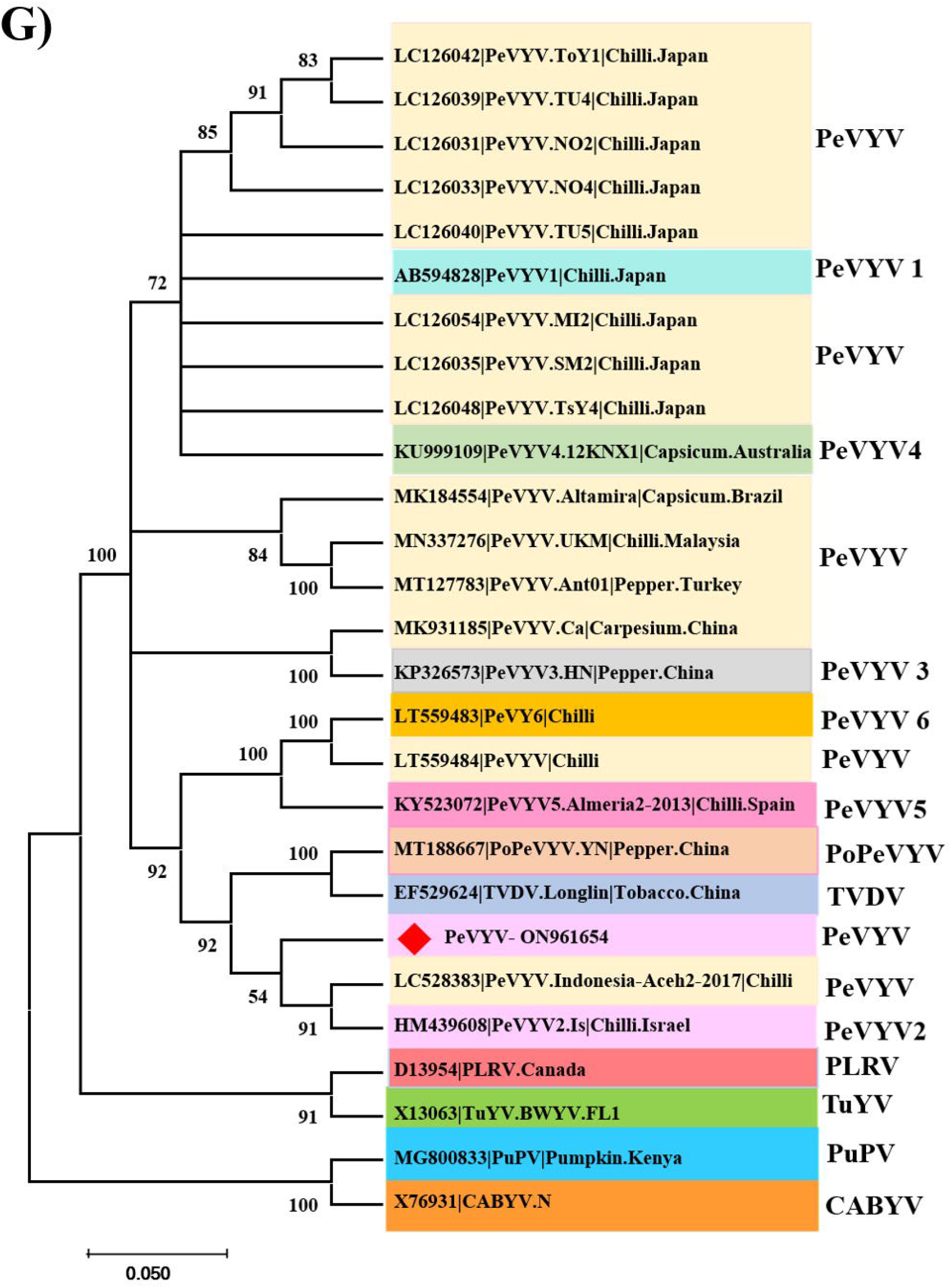
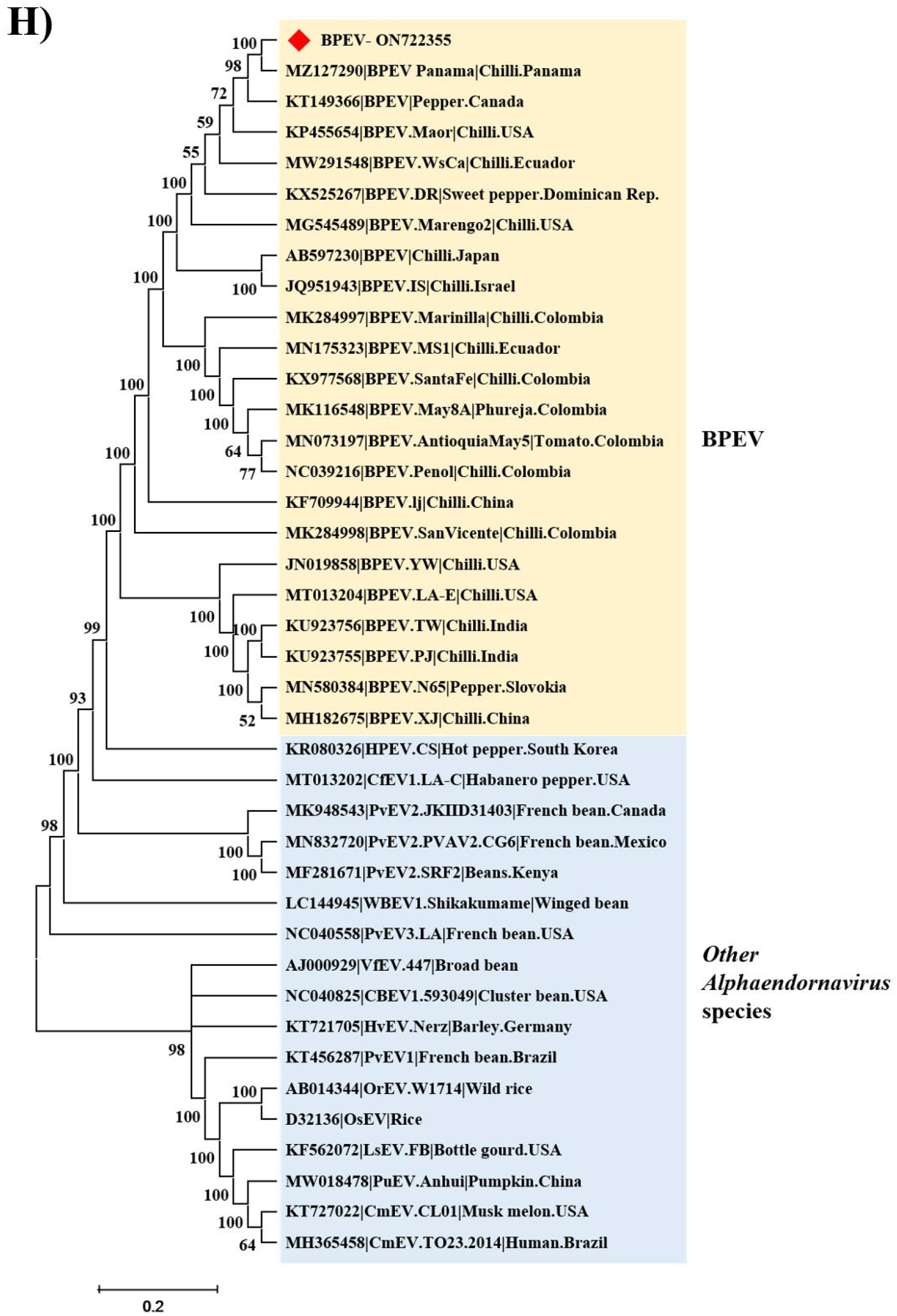
Phylogenetic tree from nucleotide sequences of reconstructed complete/near-complete viral genomes A) ChiLCV B) CMV RNA1 C) CMV RNA2 D) CMV RNA3 E) PCV-2 RNA1 F) PCV-2 RNA2 G) PeVYV and H) BPEV with other selected respective viruses using Neighbourhood Joining (NJ) method and Kimura 2-parameter (K2P) model with 1000 bootstrap replicates in MEGA 11 programme.

ChiLCV isolate (ON714166) identified in the study showed a maximum percent nt identity of 99.28% with Ahmedabad (JN663846) and Raichur (MK161454) isolates of ChiLCV and confirmed it to be an isolate of previously reported viruses (Figure 5A, supplementary table 9). Similarly, all three genome segments of CMV [RNA1 (ON722351), RNA2 (ON722352) and RNA3(ON722352)] showed close clustering with species of CMV belonging to subgroup IB (Figure 5B-D) with maximum percent nt identity with BA (MN340990; 98.13%), TNCR2-1 (MT422731; 98.11%) and KP (KM272276; 98.07%), respectively (supplementary tables 10-12). Among the two genomic segments of PCV, RNA1 (ON722354) revealed its close relation to China isolate CQ (KX905077) with maximum nt identity of 99.68% and amino acid (aa) identity of 99.22%. In case of RNA2, it showed maximum nt identity of 99.9% and 100% aa identity with South Korean isolate SC10-2 (KY923704) (Supplementary tables 13-14) and confirmed it as an isolate of PCV-2 (Figure 5E-F). PeVYV (ON961654) showed maximum percent identity with Israel isolate Is (HM439608; 93.88% nt identity and 89.54-97.78% individual genes aa identity) of PeVYV (Supplementary tables 15-16). Phylogenetic analysis showed its close clustering with isolates of PeVYV (Figure 5G). BPEV (ON722355) genome constructed under the current study, showed maximum nt identity of 99.97%with Panama isolate of BPEV (MZ127290) (Supplementary table 17). Pairwise sequence identity scores and phylogenetic analysis (Figure 5H) of the current isolate confirmed it to be a species BPEV.

### Detection of recombinants in identified viruses

Among the reconstructed viral genomes, recombination breakpoints were detected in ChiLCV (coat protein and AC4 regions), CMV RNA2 (2a protein region) and PeVYV (P0, P3 and P5 protein regions). In ChiLCV, CMV (RNA2) and PeVYV, recombination events were supported by at least three algorithms used for detection of breakpoint analysis available in the RDP5 programme. Both intraspecific and interspecific recombinants were observed in ChiLCV, CMV and PeVYV. There were no recombination breakpoints observed in CMV RNA1 and PCV-2 (RNA1 and RNA2). In remaining viral genomes like CMV (RNA3) and BPEV, recombination breakpoints were detected but the events were not supported by at least three algorithms (Supplementary table 18 and Figure 6A-C).

**Figure 6.**
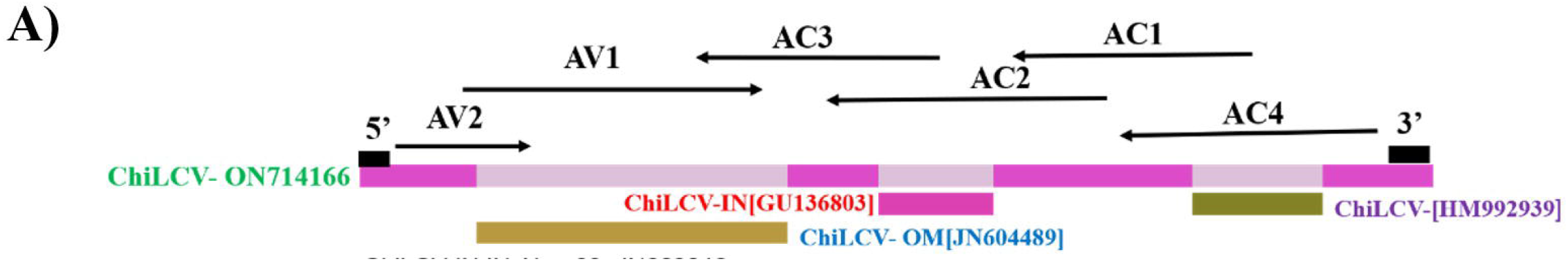
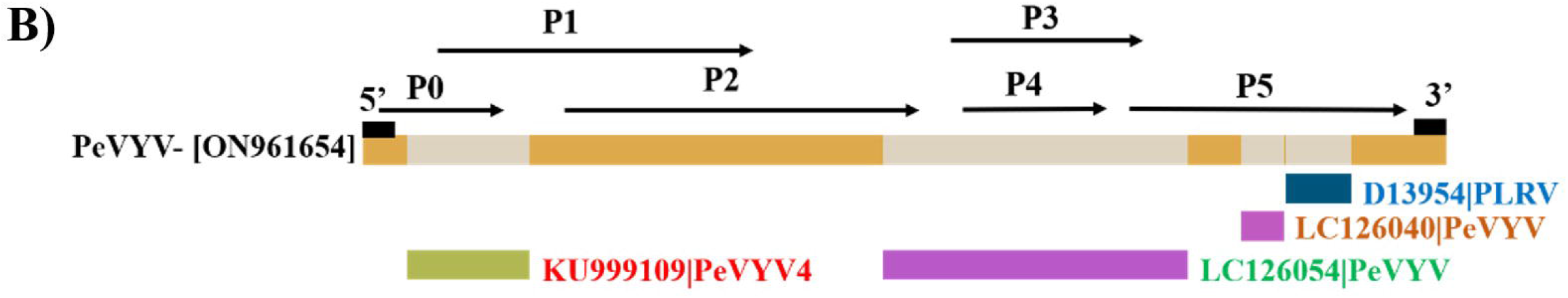
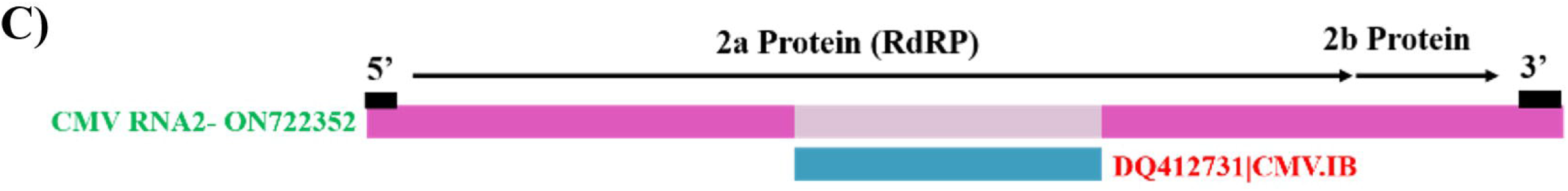
Recombination breakpoint analysis in reconstructed viral genomes. Recombination breakpoints were identified in a) ChiLCV, b) PeVYV and c) CMV RNA2. Recombinantion was observed in AV1, AV2 (coat protein) and AC4 regions of ChiLCV, P0, P3 and P5 regions of PeVYV and 2a protein (RdRP) region of CMV RNA2.

### Validation of identified viruses

Identified DNA virus (ChiLCV) was validated by PCR and LAMP. RNA viruses (CMV, GBNV, PCV-2, PeVYV and BPEV) were validated by RT-PCR and RT-LAMP using specific primers. PCR assay for DNA virus (ChiLCV) and, RT-PCR assay for RNA viruses (CMV, GBNV, PCV-2, PeVYV and BPEV) identified by NGS resulted in the expected amplicons visualized on ethidium bromide-stained 1% agarose gels (Figure 7A-F). Similarly, LAMP and RT-LAMP assays resulted in ladder-like bands further confirming the presence of identified association with chilli samples. Whereas, no amplification was observed from healthy samples and water controls in all the assays used for validation. In addition, to observe LAMP/RT-LAMP (ladder-like bands) amplified products were observed on 2% agarose gel, different dyes were used for visual detection (Figure 8A-F) without going for electrophoresis.

**Figure 7.**
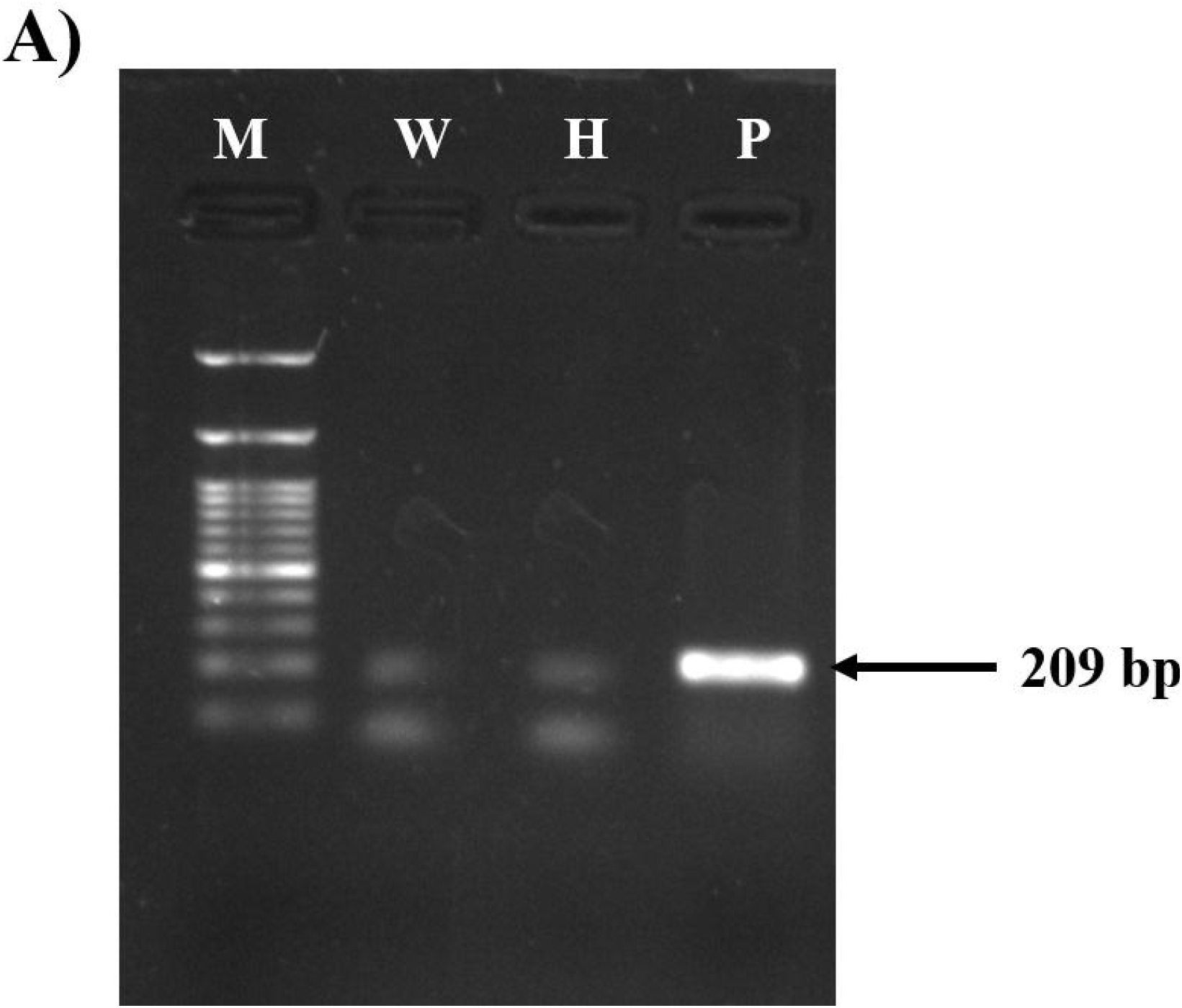
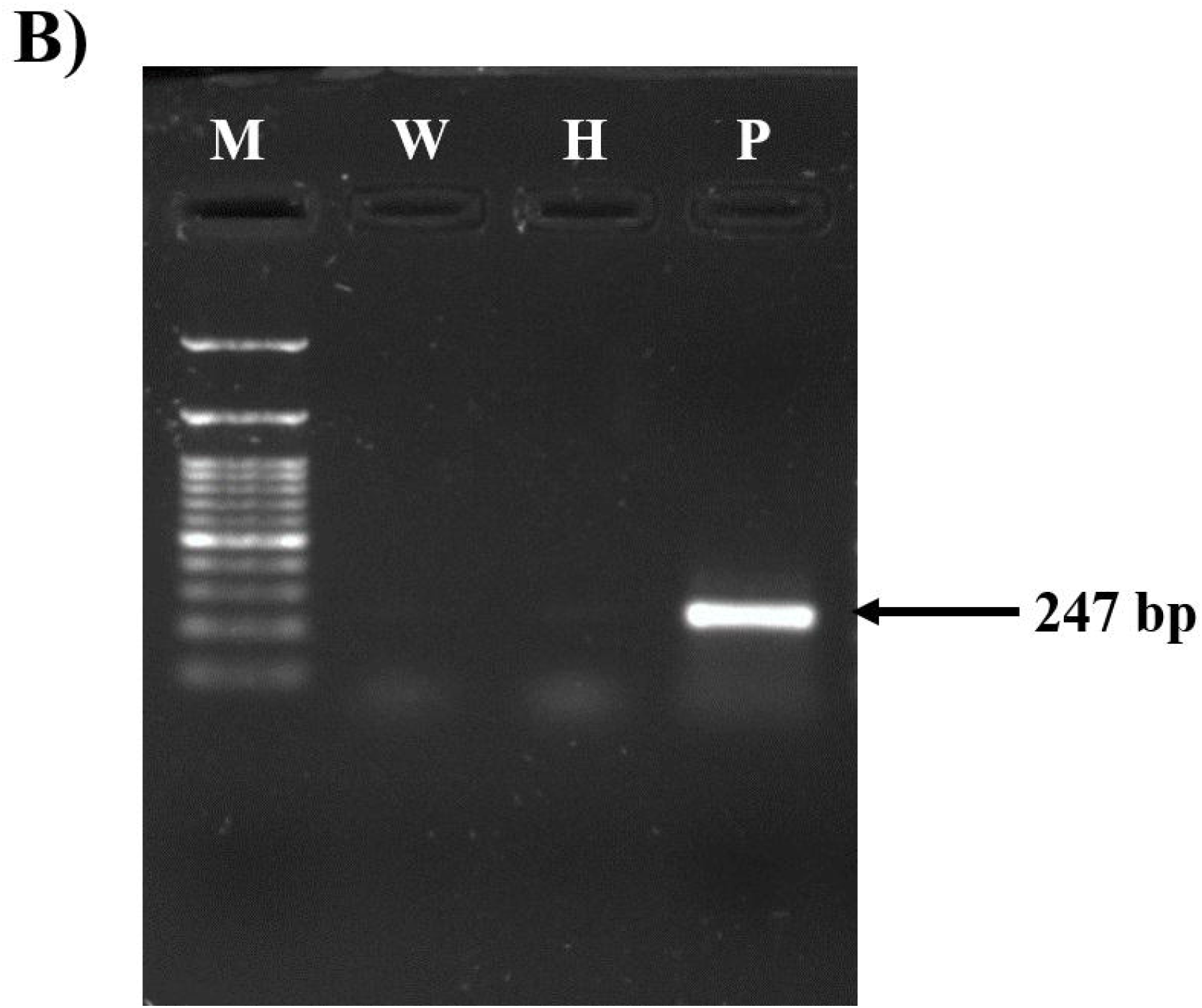
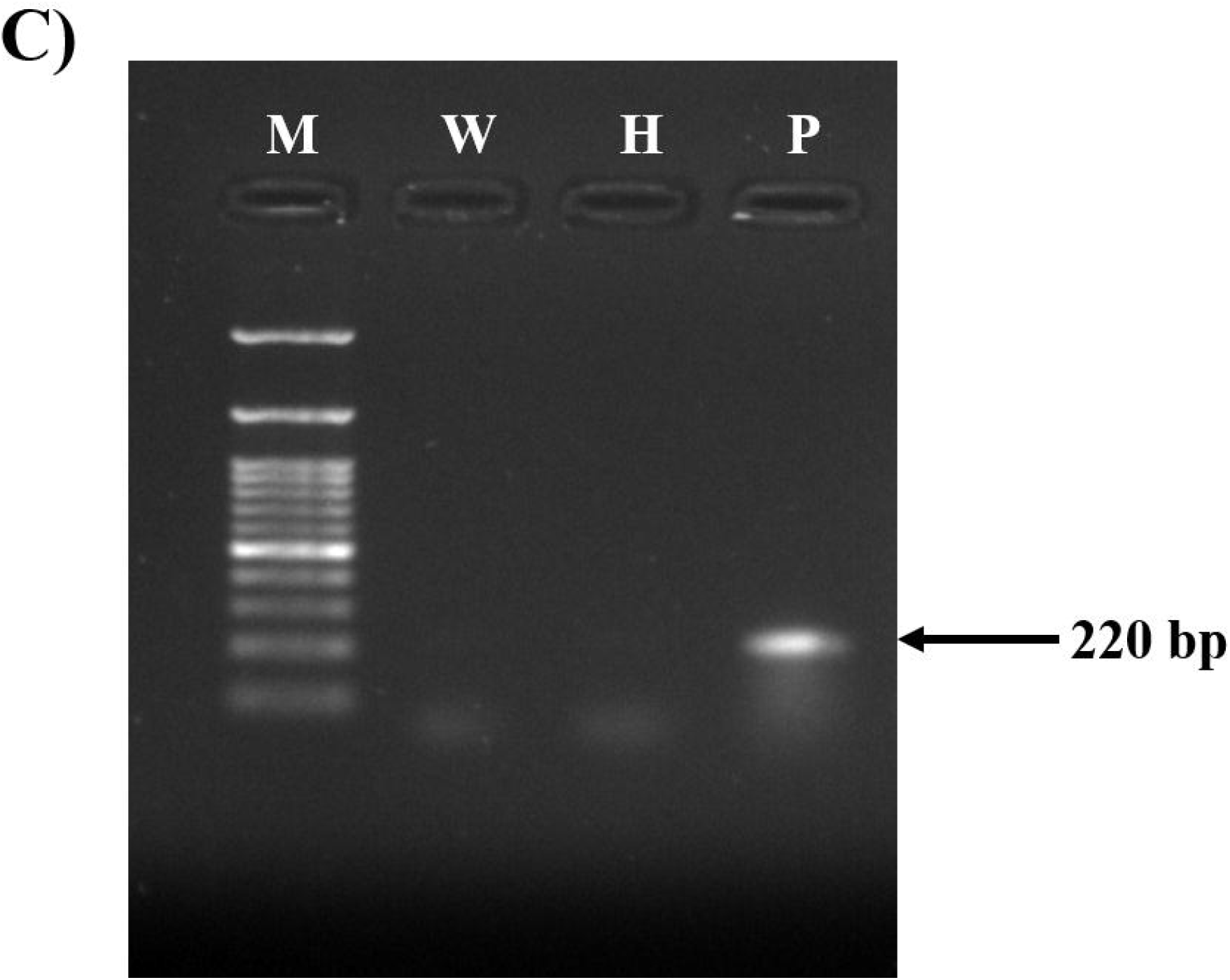
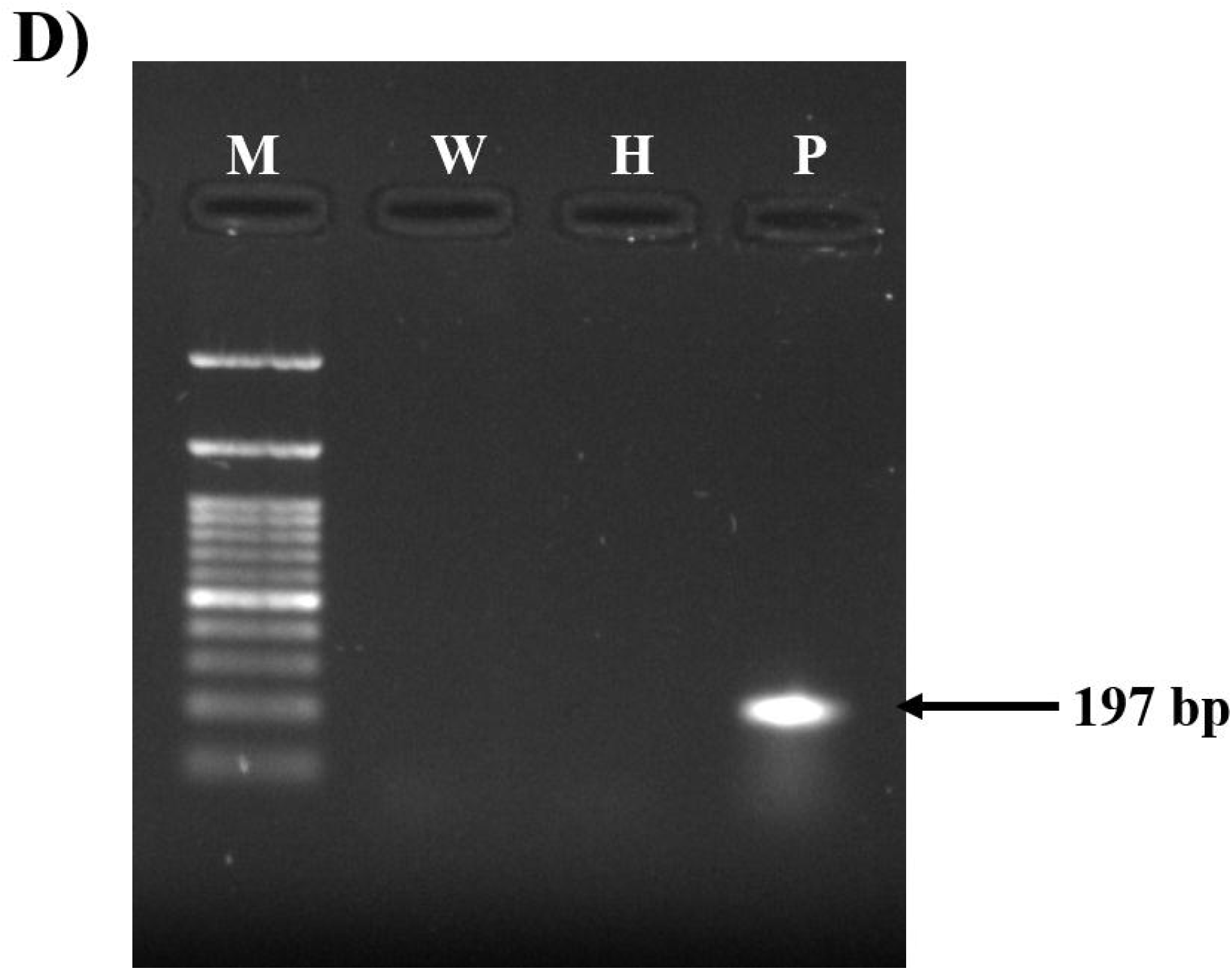
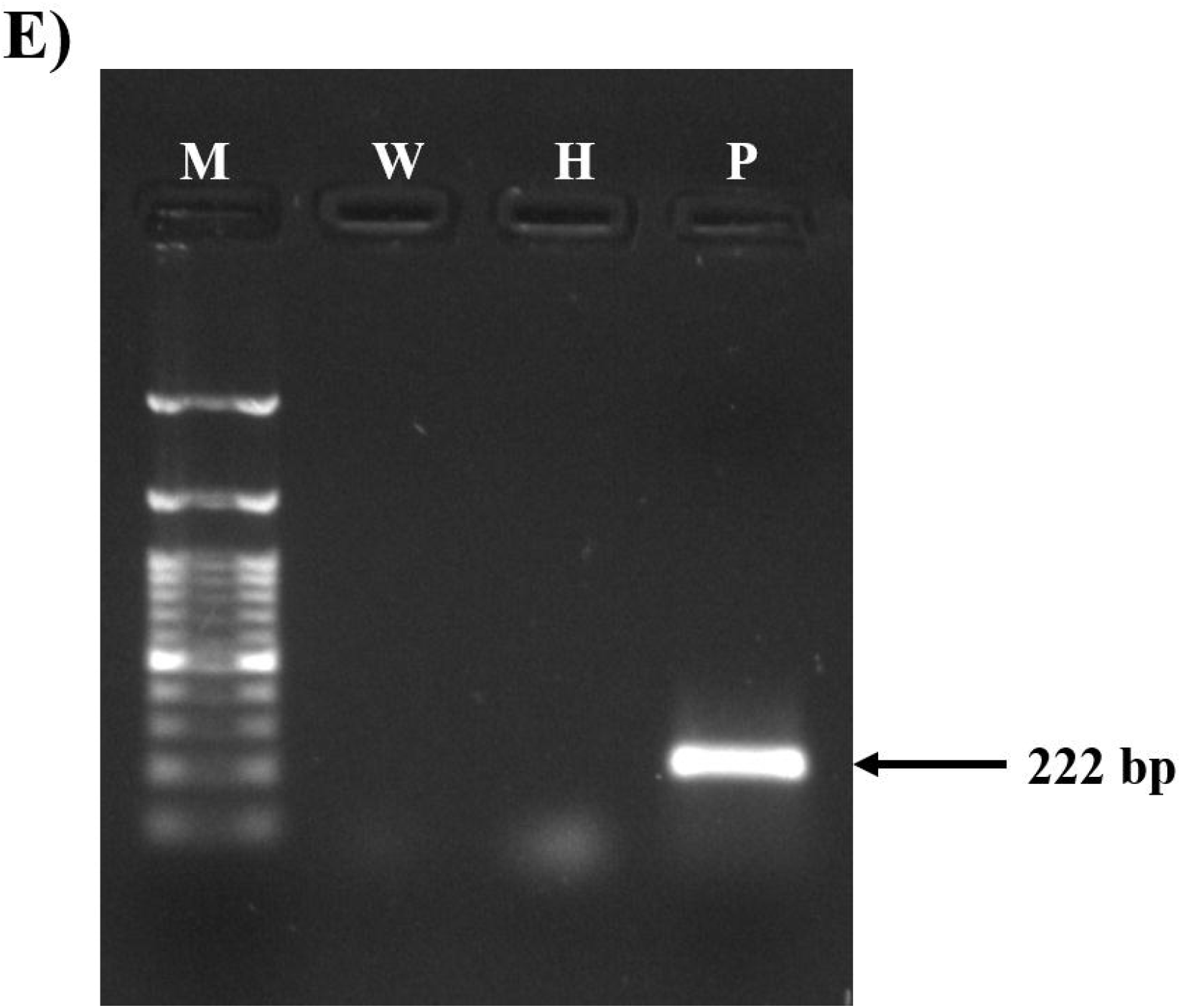
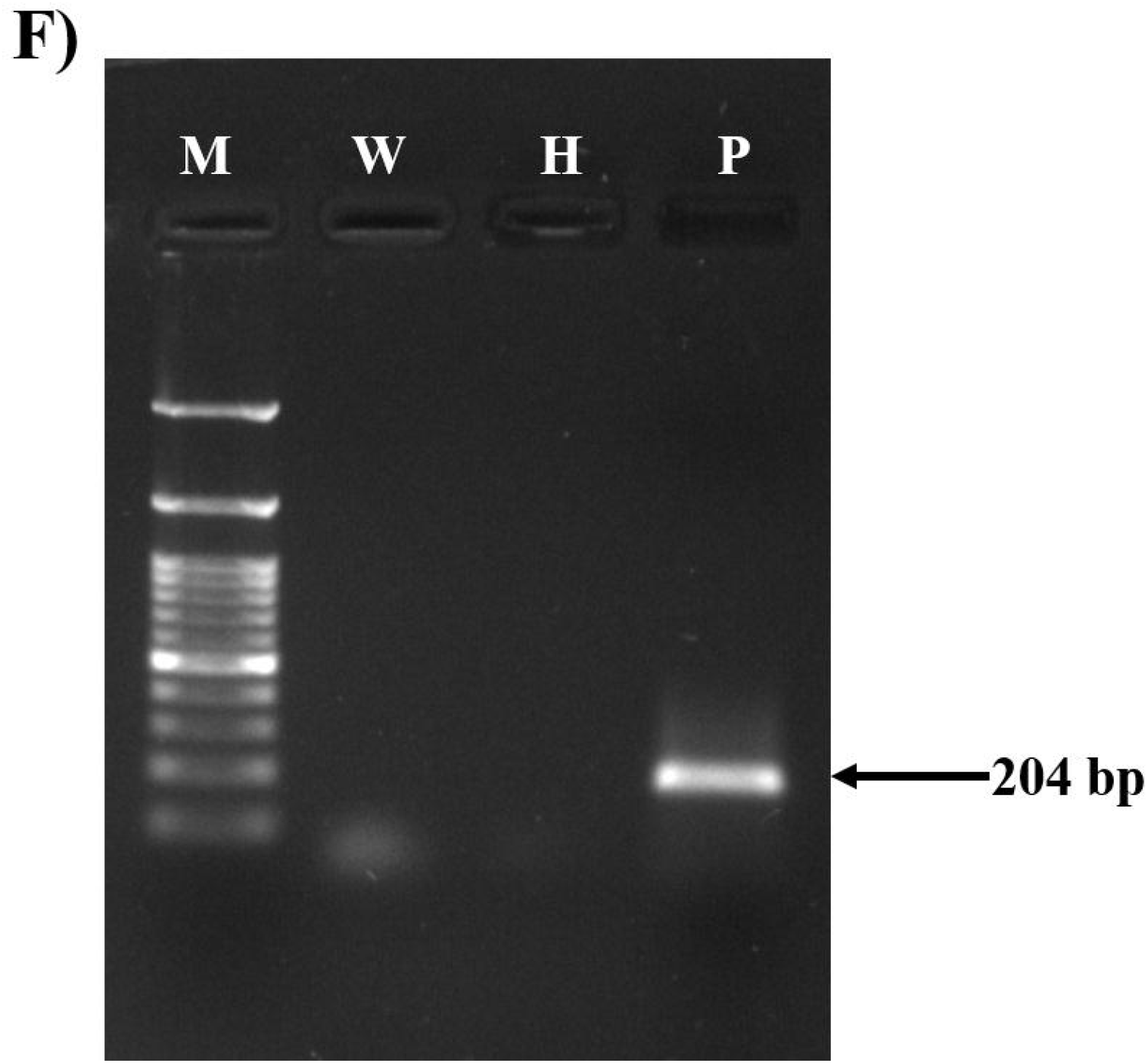
Validation of identified viruses A) ChiLCV with PCR and B) CMV, C) GBNV, D) PCV-2, E) PeVYV and F) BPEV with RT-PCR assay. PCR/RT-PCR amplification products were checked on 1% agarose gel. Lane M: 100 bp DNA ladder, Lane W: Water control, Lane H: Healthy sample and Lane P: Pooled DNA/RNA sample used for virome analyses. Coat protein gene was amplified in ChiLCV, CMV, PCV-2 and PeVYV, movement protein gene was amplified in GBNV and polyprotein gene was amplified in BPEV.

**Figure 8.**
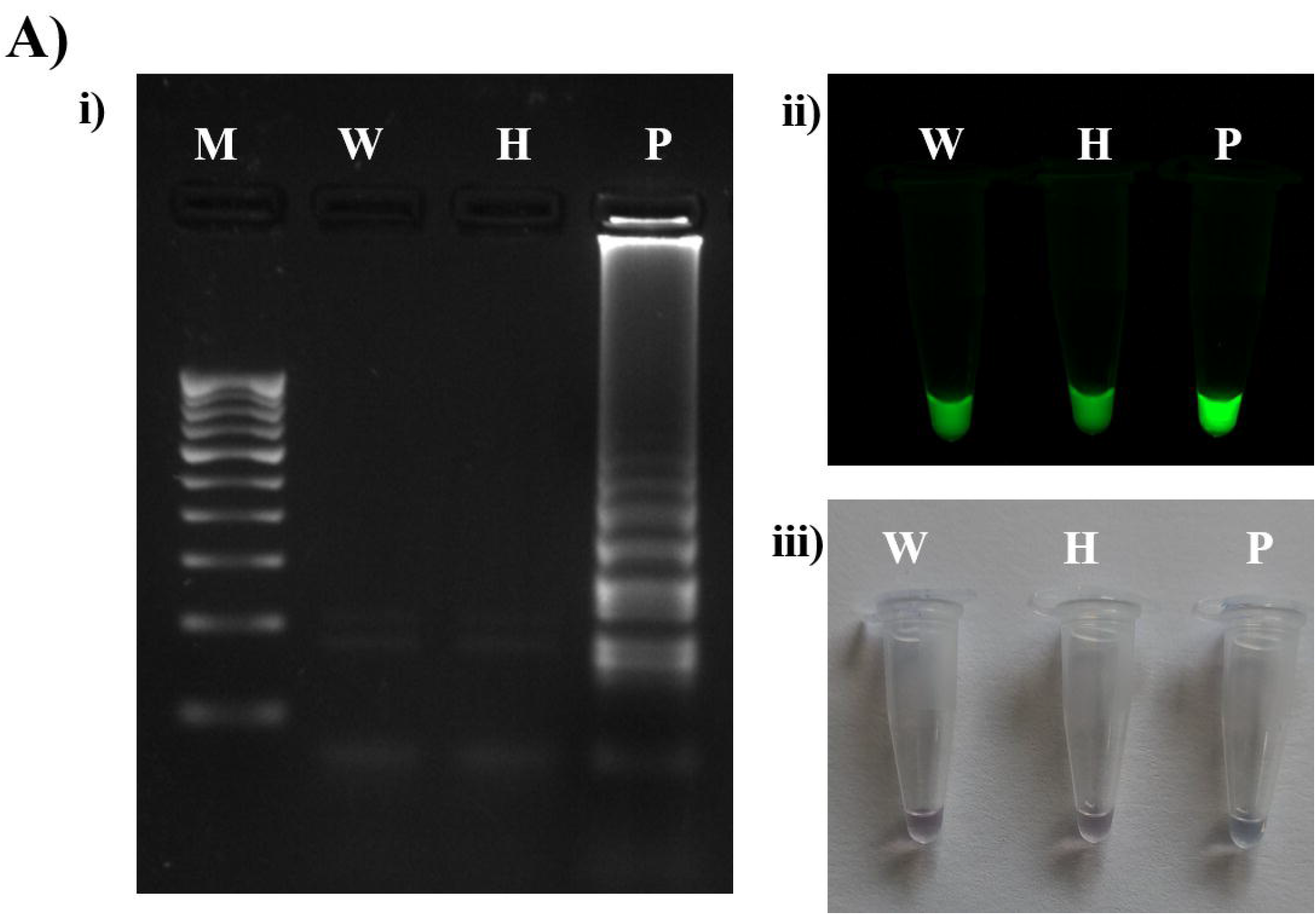
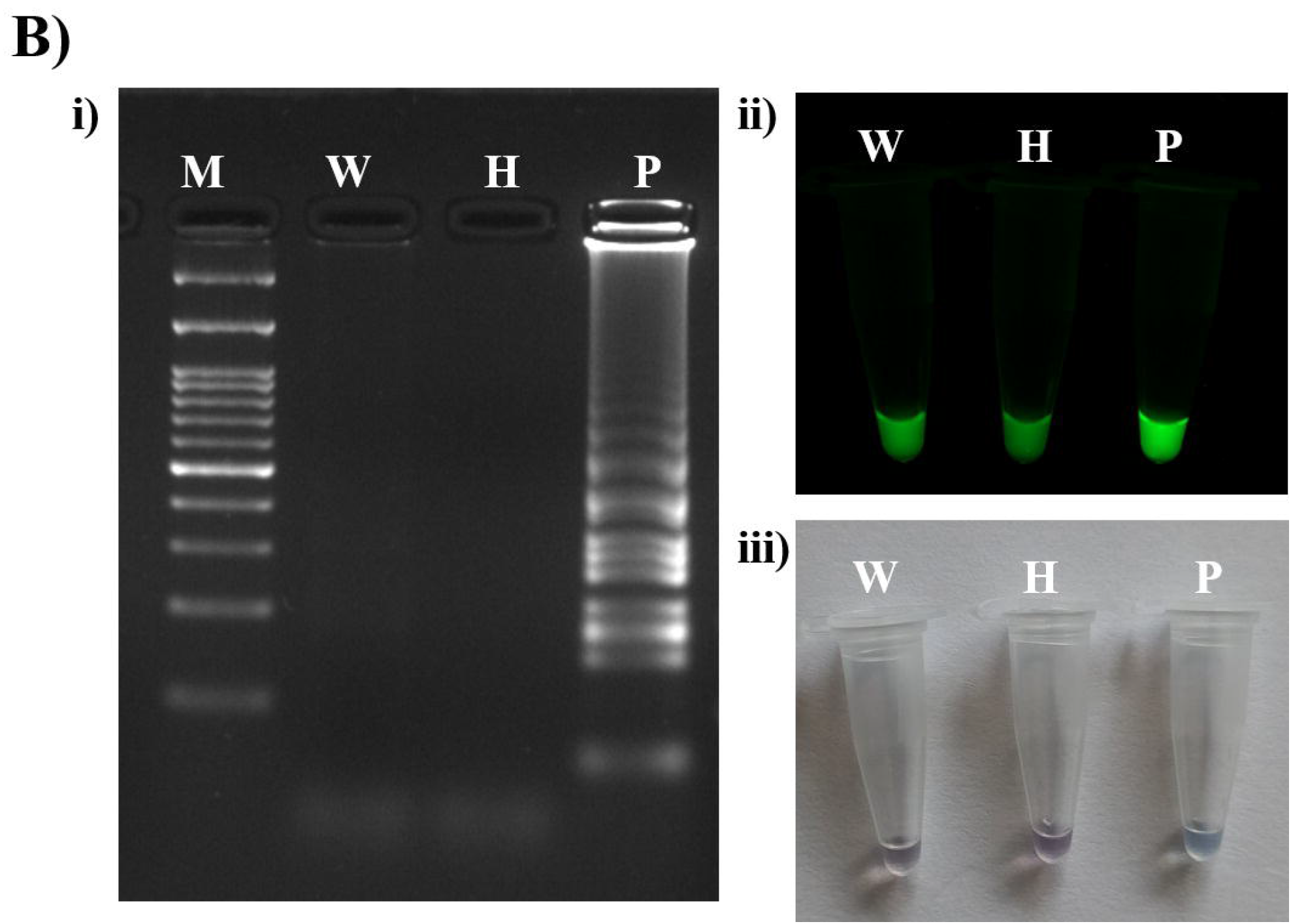
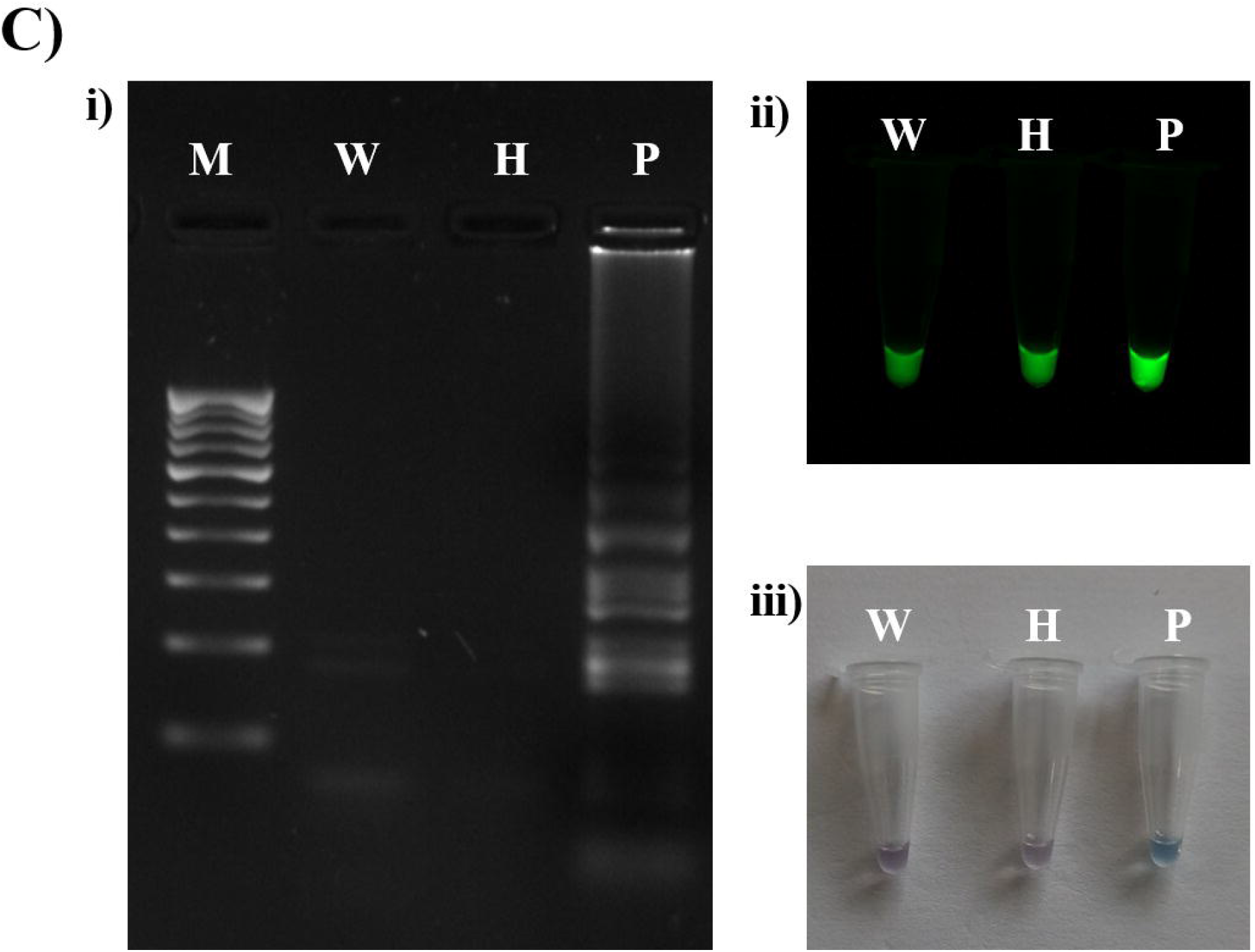
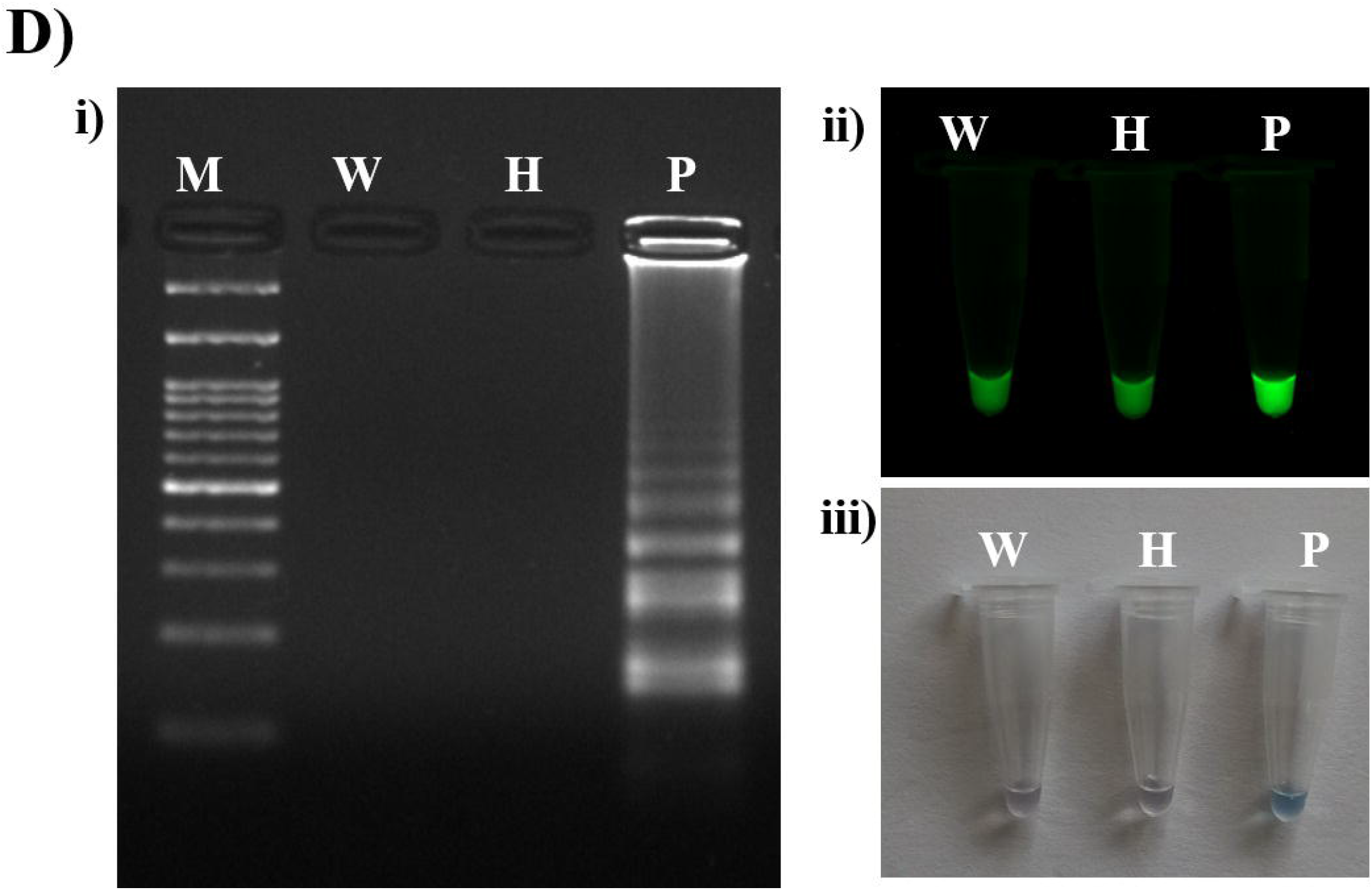
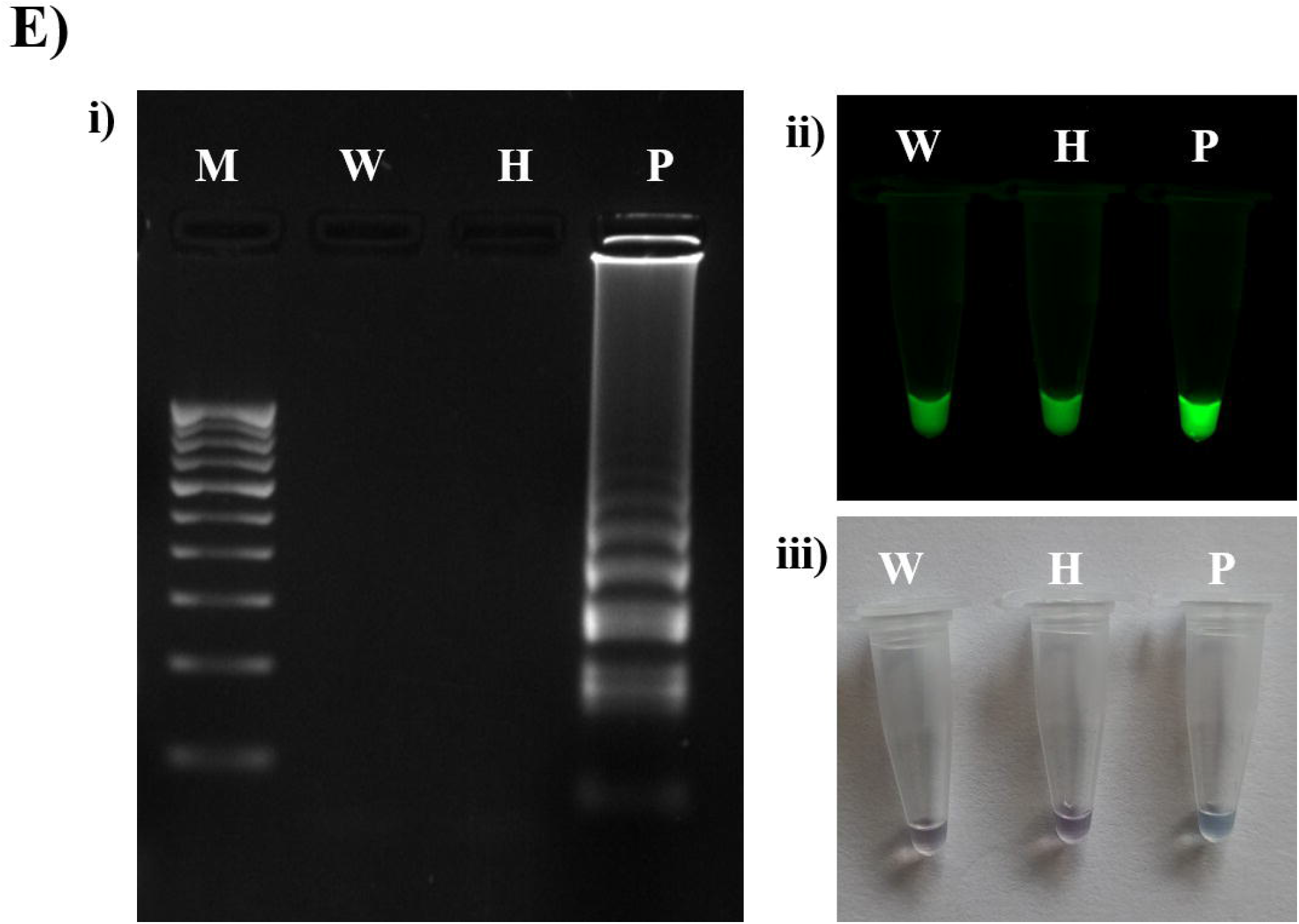
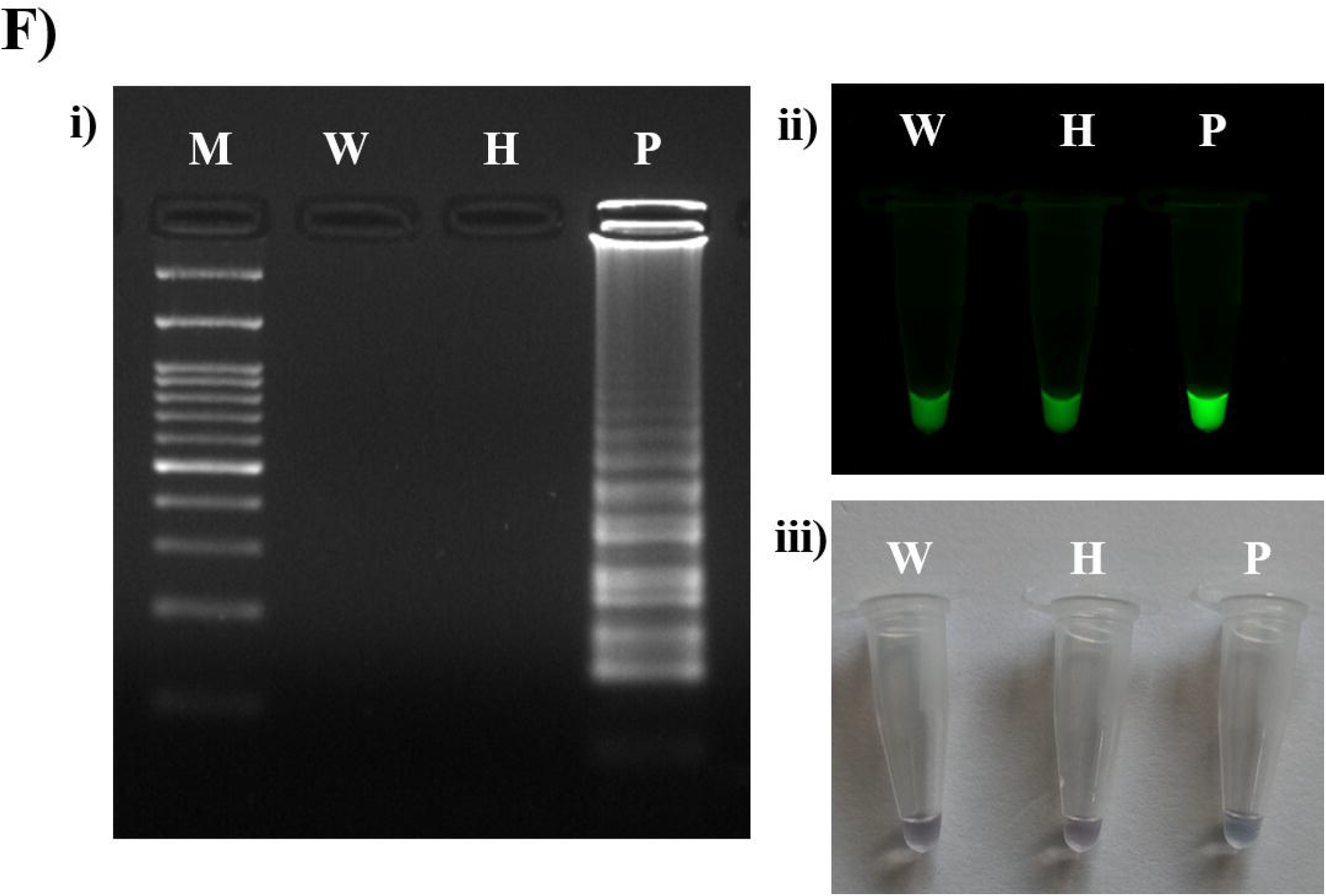
Validation of identified viruses A) ChiLCV with LAMP and B) CMV, C) GBNV, D) PCV-2, E) PeVYV, F) BPEV with RT-LAMP assays. LAMP/RT-LAMP reaction products on 2% agarose gel ii) Visual inspection with Veri PCR dye. Pooled DNA/RNA sample used for virome analyses showed green fluorescence, whereas in healthy sample and water control no fluorescence was observed. iii) Visual inspection with hydroxy naphthol blue (HNB). Pooled DNA/RNA sample used for virome analyses showed change in colour from violet to sky blue, whereas in healthy sample and water control no change in violet colour was observed. Lane M: 100 bp DNA ladder, Lane W: Water control, Lane H: Healthy sample and Lane P: Pooled DNA/RNA sample used for virome analyses. Coat protein gene was amplified in ChiLCV, CMV, PCV-2 and PeVYV, movement protein gene was amplified in GBNV and polyprotein gene was amplified in BPEV.

## DISCUSSION

Chilli is one of the major commercial spice and vegetable crops grown across the world. Several viral diseases are the constraints for its production and resulting in huge economic losses to the farmers. Many viruses causing the disease were identified in chilli using traditional techniques (ELISA and PCR-based techniques) (Baruah et al., 2016). Diagnosis of plant virus diseases by traditional methods is carried out for the purpose of their prevalence, identifying the host range, virus association with vectors, virus indexing in seed/planting material, quarantine regulations, screening of breeding material etc. (Rubio et al., 2020). By following the traditional diagnostic techniques, only known viruses can be detected leaving behind unknown viruses, which is true in chilli as well. Advancements in the sequencing arena using the NGS platforms, enabled researchers to overcome the limitations of traditional methods in identifying unknown viruses (Hadidi et al., 2016). NGS is gaining importance and becoming most sought tool for detection and identification of known and novel viruses (Pecman et al., 2017).

In the current study, an attempt was made for virome profiling of chilli leaf samples showing different kinds of disease symptoms such as leaf curl, vein banding, mosaic, mottling, shoestring/rat tail/filiform/leathery and dull colour. According to our knowledge, this is the first virome analyses study in chilli from India to identify known and novel viruses. In nature, it is not surprising to detect mixed infection in a plant (Moreno and López-Moya, 2020), and it results in varied symptoms due to dominance, masking or synergism of viruses. Interaction/synergism between plant viruses in host might lead to unpredictable variations in symptomatology, infectivity and transmissibility (Martín and Elena, 2009). Survey in the major chilli growing districts (Kolar and Bengaluru) of Karnataka State, India revealed the ubiquitous prevalence of viral diseases with diverse symptoms and incidences ranging from 26.66% to 47.50%. Nineteen leaf samples showing difference with symptom phenotype were collected and virome profiling was done. Total RNA extracted from each sample was pooled at an equimolar concentration to get a representative RNA pool for all samples and data on mRNAome and sRNAome was obtained by sequencing the rRNA depleted pooled total RNA. Similar approach was followed by many researchers in unravelling the viruses infecting agricultural and horticultural crops (Seguin et al., 2014; Roossinck et al., 2015; Sidharthan et al., 2020). Hence, it might be ideal to use both ribosomal RNA depleted libraries (mRNAome and sRNAome) for studies focusing on detection of viruses (Jo et al., 2015).

Both mRNAome, sRNAome and whole transcriptome (WT) were assembled *de novo* due to unavailability of chilli reference genome. The *de novo* assembly of transcripts depends on the information contained in the reads alone and these assemblies have an added benefit of identifying novel transcript isoforms and alternative splicing events (Martin and Wang, 2011; Zhao et al., 2011; Wong et al., 2018). In recent studies, viral genomes have been recovered from *de novo* assembled mRNA and sRNA (Kreuze et al., 2009; Jo et al., 2015). In general, paired-end reads have more information than single reads (Isnard et al., 1998). In this study, the paired-end reads were *de-novo* assembled by following three approaches, mRNAome with Trinity, sRNAome with Velvet and WT with SPAdes *de novo* assemblers. Differential utilization of these *de novo* assembling tools was well proven to yield better results in several studies (Hölzer and Marz, 2019; Sidharthan et al., 2020). Further, when multiple assemblers were used, it will help in better identification of viruses in chilli virome eliminating the chances of not detecting while using a single assembler (Massart et al., 2019). Trinity assembly with mRNA and SPAdes assembly with WT libraries yielded longer contigs compared to Velvet assembly with sRNA library. However, the shorter reads/contigs from sRNAome can be useful for identification of viruses with low titres (Czotter et al., 2018).

Seven viruses in the chilli samples were identified by using Trinity and SPAdes assemblers and five viruses by Velvet assembler. From all three assemblies we identified seven viruses (ChiLCV, CMV, GBNV, PCV-2, PeVYV, BPEV and TVCV), five satellites of DNA viruses (ToLCBDB, ToLCAA, CYVMA, AYVSA and ToLCVirA) found to be associated with chilli samples presented with the diverse phenotypic symptoms. All five satellite viruses were identified from sRNA assembly in comparison to mRNAome and WT, where, only three of the DNA satellites of viruses (ToLCBDB, CYVMA and ToLCVirA) were identified. This might be due to the more number of shorter contigs represented by sRNAome than the mRNAome and WT (Maliogka et al., 2018).

Length of the viral contig is an important factor for identifying viruses through NGS data. Because the shorter contigs might reveal the presence of various viruses belonging to same genus although these contigs could be derived from a single virus genome. Therefore, the BLAST results sometimes lead to a mismatch in virus identification due to shorter viral fragments and it is advised to use complete or nearly complete viral genomes sequences for identification of viral genomes (Jo et al., 2017). Complete or near complete viral genomes were constructed using virus associated contigs extracted separately from mRNAome, sRNAome and WT data. Further contigs were mapped to their respective reference sequences present in NCBI database. This resulted in construction of complete or nearly complete genomes of five viruses: ChiLCV, CMV, PCV-2, PeVYV and BPEV and partial genome for two viruses: GBNV and TVCV. Incomplete recovery of viruses could be due to presence of lower virus titre, silencing of virus by host, poor RNA quality, sequencing depth and analysis of the data (Thakur and Varshney, 2010). Even though ChiLCV is a DNA virus, the whole genome can be constructed from mRNA transcriptome data, which covers most regions of virus-encoded genes. Jo et al. (2017) was successful in recovering geminiviruses and two satellite DNAs from pepper virome obtained from mRNA data.

Further, for sequence comparison and phylogenetic analysis complete/nearly complete genomes were used. Based on the current species demarcation criteria for begomoviruses (91% nucleotide sequence identity) (Walker et al., 2021), ChiLCV (ON714166) recovered from the current study, confirmed it be a variant of ChiLCV isolates of Ahmedabad and Raichur. In case of CMV (RNA1, RNA2 & RNA3), at least 65 per cent of whole genome sequence identity (Bujarski, 2021) should match with the deposited sequences in NCBI database. Three RNA segments constructed in the study are closely related to CMV subgroup IB. The phylogenetic analysis of PCV recovered from chilli samples revealed its closeness to the species of PCV-2 and also met the criteria for species demarcation (90 and 80 per cent aa identity in the RNA1 (RdRP) and RNA2 (CP), respectively) (Vainio et al., 2018) of deltapartitiviruses. If a virus is to be assigned as species of poleroviruses, then differences in aa sequence of any gene product should not be greater than 10% (Sõmera et al., 2021). These criteria were fulfilled by the genome of PeVYV recovered from chilli virome in the current study. BPEV constructed from assembled data showed a maximum nt identity of 99.97% with Panama isolate of BPEV and considered as its isolate based on the species demarcation criteria for alphaendornaviruses (75% whole genome sequence identity) (Valverde et al., 2019).

Recombination is one of the important factors for emergence of viruses through different means of recombination and plays a crucial role in emergence of novel viruses (Seal et al., 2006). Unlike sexual reproduction, recombination in viruses involves the exchange/transfer of genomic fragments between the members of distantly related species of viruses. This might create a novel recombinant genome with an added advantage of adaptivity to a hostile environment, expansion of host range and production of new symptoms (Martin et al., 2011). Recombination analysis revealed presence of recombination breakpoints in ChiLCV (coat protein and AC4 regions). Similarly, recombination breakpoints were detected in the P0, P3 and P5 protein regions of PeVYV and CMV RNA2 (2a protein region).

Validation for the identified viruses in the pooled nucleic acid (DNA/RNA) samples extracted from infected chilli leaf samples was done by two methods, PCR and LAMP for DNA virus (ChiLCV) and RT-PCR and RT -LAMP for RNA viruses (CMV, GBNV, PCV-2, PeVYV and BPEV). Presence of all the viruses identified in the pooled RNA extracted from nineteen samples using NGS was validated by PCR and LAMP variant detection protocols. According to our knowledge, this is the first time LAMP assay was used for the detection of ChiLCV, GBNV, PCV-2, PeVYV and BPEV in India. Whereas, LAMP assay for detection of CMV was previously done in pepper by Bhat et al. (2013). As these LAMP/RT-LAMP detection of plant viruses is highly specific and sensitive over PCR and RT-PCR (Saharan et al., 2013; Wong et al., 2018; Soroka et al., 2021). The confirmation of identified viruses with molecular detection methods will strengthen the studies of virome analyses by NGS.

The current study concludes that 1). chilli plants showing viral disease symptoms in Bengaluru and Kolar Districts of Karnataka State, India are associated with seven different viruses (ChiLCV, CMV, GBNV, PCV-2, TVCV, PeVYV and BPEV) and five DNA satellites (ToLCBDB, ToLCAA, CYVMA, AYVSA and ToLCVirA). 2). PCR and LAMP based tools developed by designing the primers for identified viruses can be utilized for their routine detection protocols 3). This is the first report of two viruses, PeVYV and BPEV associated with any crop plant with disease phenotype in India. Information reported here provides a snapshot of chilli virome and adds new knowledge about viruses associated with chilli.

## DATA AVAILABILITY STATEMENT

The datasets analyzed in this study are available in the NCBI repository under the Bioproject PRJNA914908 with accession numbers, SRR22863580 and SRR22876797, respectively for mRNA and sRNA. The complete or near-complete viral genomes: ChiLCV (ON714166); CMV-RNA1 (ON722351), RNA2 (ON722352), RNA3 (ON722352), PCV-2-RNA1 (ON722354) and RNA2 (ON645206), PeVYV (ON961654) and BPEV (ON722355) have been submitted to NCBI GenBank.

## AUTHOR CONTRIBUTIONS

NVR generated the samples for sequencing and done the validation work, NVR, SH and PK performed bioinformatic analyses. MM, HDV, KSS, VV and CRJB performed diversity analysis. CNLR conceptualized work and provided the overall direction. All the authors are involved in literature mining and manuscript preparation. All authors read and approved the final manuscript.

## TABLE LEGENDS

## FIGURE LEGENDS

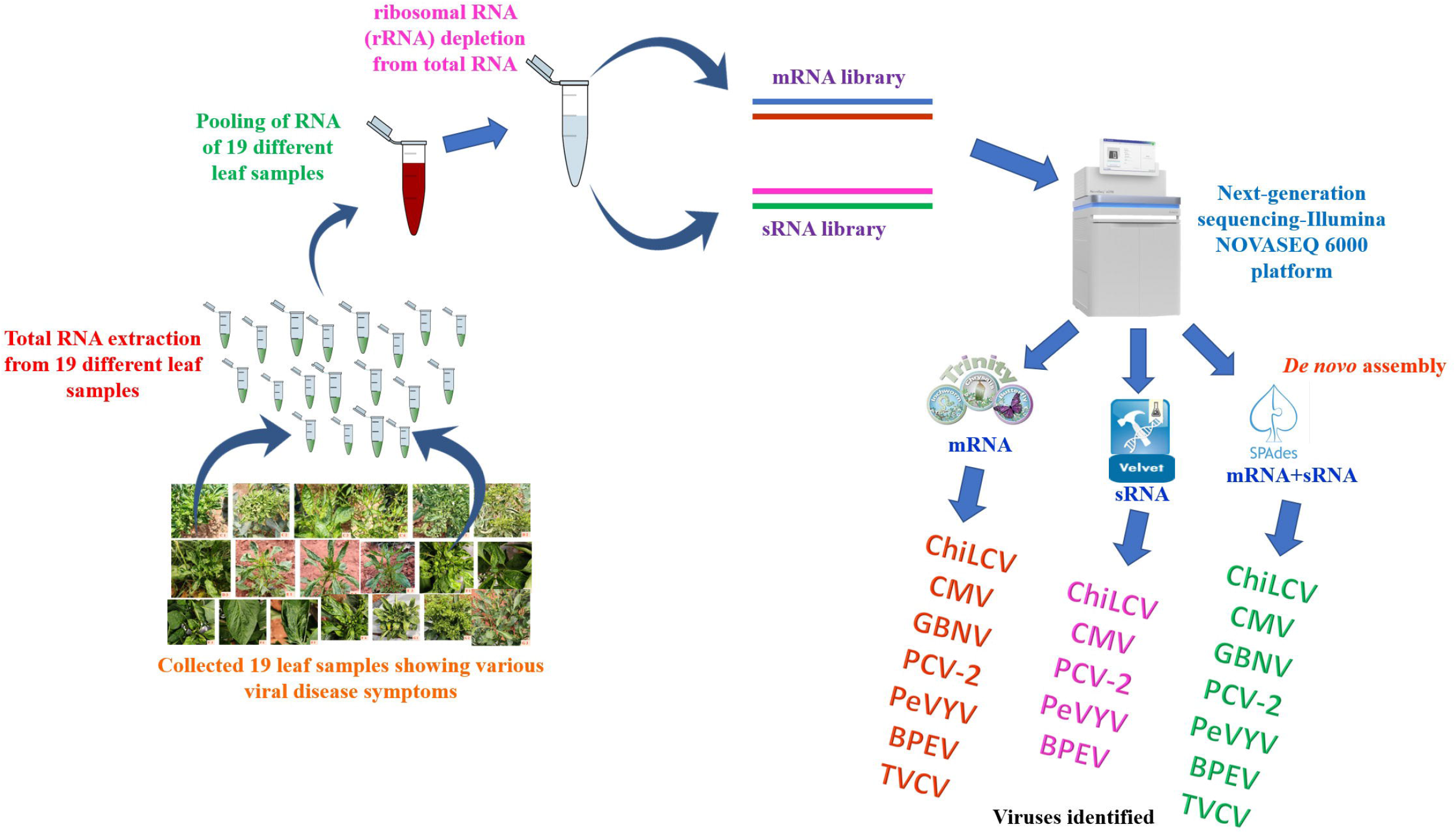

